# Mesocorticolimbic circuit mechanisms underlying the effects of ketamine on dopamine: a translational imaging study

**DOI:** 10.1101/748665

**Authors:** Michelle Kokkinou, Elaine E. Irvine, David R. Bonsall, Sridhar Natesan, Lisa A. Wells, Mark Smith, Justyna Glegola, Eleanor J. Paul, Kyoko Tossell, Mattia Veronese, Sanjay Khadayate, Nina Dedic, Seth C. Hopkins, Mark A. Ungless, Dominic J. Withers, Oliver D. Howes

## Abstract

Patients with schizophrenia show increased striatal dopamine synthesis capacity in imaging studies. However, the mechanism underlying this is unclear but may be due to N-methyl-D-aspartate receptor (NMDAR) hypofunction and parvalbumin (PV) neuronal dysfunction leading to disinhibition of mesostriatal dopamine neurons. Here, we test this in a translational mouse imaging study using a ketamine model. Mice were treated with sub-chronic ketamine (30mg/kg) or saline followed by *in-vivo* positron emission tomography of striatal dopamine synthesis capacity, analogous to measures used in patients. Locomotor activity was measured using the open field test. *In-vivo* cell-type-specific chemogenetic approaches and pharmacological interventions were used to manipulate neuronal excitability. Immunohistochemistry and RNA sequencing were used to investigate molecular mechanisms. Sub-chronic ketamine increased striatal dopamine synthesis capacity (Cohen’s d=2.5, *P<0.001)* and locomotor activity. These effects were countered by inhibition of midbrain dopamine neurons, and by activation of cortical and ventral subiculum PV interneurons. Sub-chronic ketamine reduced PV expression in these neurons. Pharmacological intervention with SEP-363856, a novel psychotropic agent with agonism at trace amine receptor 1 (TAAR1), significantly reduced the ketamine-induced increase in dopamine synthesis capacity. These results show that sub-chronic ketamine treatment in mice mimics the dopaminergic alterations in patients with psychosis, and suggest an underlying neurocircuit involving PV interneuron hypofunction in frontal cortex and hippocampus as well as activation of midbrain dopamine neurons. A novel TAAR1 agonist reversed the dopaminergic alterations suggesting a therapeutic mechanism for targeting presynaptic dopamine dysfunction in patients.

## INTRODUCTION

Schizophrenia is a severe mental disorder and a significant global health burden, highlighting the need to better understand its neurobiology in order to develop improved treatments (1). Dopaminergic hyperactivity in the striatum is thought to underlie the symptoms of schizophrenia, particularly psychosis (2–5). Supporting this, 3,4-dihydroxy-6-^18^F-fluoro-l-phenylalanine ([^18^F]-FDOPA) positron emission tomography (PET) imaging studies have revealed higher striatal dopamine synthesis capacity in patients with schizophrenia (6–9). Furthermore, increased dopamine synthesis capacity is associated with both the development of psychosis (10) and the severity of symptoms (11). Currently available antipsychotics are all dopamine receptor blockers, which are inadequate and poorly tolerated in many patients, and do not address the mechanism underlying the dopamine dysfunction (12, 13). In addition to dopaminergic dysfunction, the glutamate hypothesis of schizophrenia has developed from the observations that N-methyl-D-aspartate receptor (NMDAR) antagonists such as ketamine induce psychotic symptoms in healthy humans and exacerbate symptoms in patients (14, 15). Furthermore, schizophrenia is associated with a reduction in parvalbumin (PV)-expressing GABAergic interneurons, which are regulated by NMDAR in the cortex and hippocampus (16–18). It has been suggested that impaired PV neuronal function in the cortex and hippocampus may lead to disinhibition of mesostriatal dopamine neuron activity via a polysynaptic pathway (19). However, the circuit mechanisms underlying increased dopamine synthesis capacity in patients, and how these are regulated by changes in PV neuronal functions, remain largely unknown.

To address this, we tested the effect of sub-chronic ketamine administration on dopamine synthesis capacity in mice using the same [^18^F]-FDOPA PET imaging technique that previously demonstrated elevation in dopamine synthesis capacity in patients (6–9). Subsequently, we investigated the underlying neurocircuit alterations using cell-type-specific chemogenetic manipulations. In order to gain insights to the translational potential of such approaches, dopamine synthesis capacity was also assessed following treatment with SEP-0363856 (SEP856), a novel psychotropic agent that inhibits ventral tegmental area (VTA) neuronal firing possibly through agonism of the trace amine receptor 1 (TAAR1) (20). Our objective was to develop a chemogenetics/PET approach that is translationally relevant and provides novel insights into the pathophysiology of schizophrenia.

## METHODS AND MATERIALS

All experiments were approved by the UK Home Office under the Animal (Scientific Procedures) Act (ASPA) 1986 and Regulation 7 of the Genetically Modified Organisms (Contained Use) Regulations 2000. All procedures were performed in accordance with the ASPA 1986 and EU directive 2010/63/EU as well as being approved by Imperial College Animal Welfare and Ethical Review Body.

### Subjects

Male mice were 6-8 weeks of age at the time of stereotaxic surgeries and 8-10 weeks of age at the start of the experiments. C57BL/6 wild-type, Dopamine transporter (DAT) Cre (*DAT∷Cre*) and Parvalbumin (PV) Cre (*PV∷Cre*) mice maintained on a C57BL/6 background were used.

### Sub-chronic ketamine regime

Ketamine hydrochloride solid (Sigma Aldrich) was dissolved in 0.9% saline solution to 6mg/ml and injected at a volume of 5ml/kg of body weight, thus administered at a dose of 30mg/kg (i.p) once daily for five consecutive days (Figure 1a, Supplementary Figure 1a, Supplementary Figure 2a, Supplementary Figure 3a) (21). Control mice received an equivalent volume of 0.9% saline vehicle.

**Figure 1:**
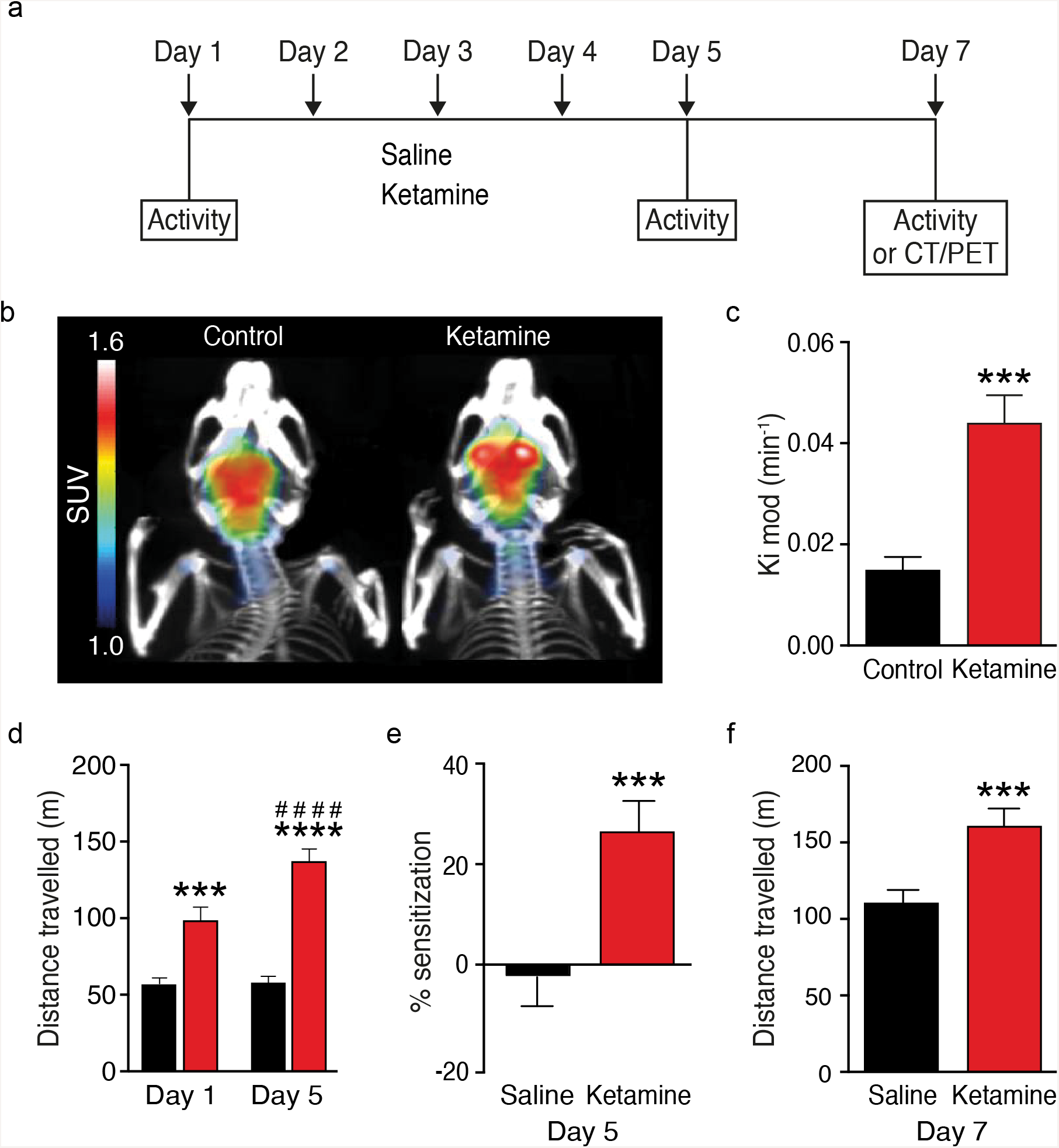
Sub-chronic ketamine increases dopamine synthesis capacity and locomotor activity. (a) Schematic showing the drug treatment schedule used to study the effect of sub-chronic ketamine administration on striatal presynaptic dopamine synthesis capacity and locomotor activity in the mouse. (b) [^18^F]-FDOPA PET brain image (averaged from 20 to 90 minutes) demonstrating high signal to noise ratio specificity in striatal uptake coregistered to the CT mouse brain image from mice treated with saline or ketamine. Standardized uptake value (SUV) activity is presented as summed activity over the timeframe (20min-90min) used to measure dopamine synthesis capacity (indexed as the uptake rate constant *K*_i_ ^mod^). (c) Striatal dopamine synthesis capacity (*K*_i_ ^mod^/minute) is significantly increased in the ketamine-treated (n=8) group versus control (*n*=7) group (\*\*\**P<0.001*, two-tailed, Cohen’s d=2.5, t_13_= 4.74)). (d) Total distance travelled during 30min post drug administration. There was a significant effect of group (F _(1, 26)_ = 46.21, *P<0.0001*), day (F _(2, 52)_ = 23.27, *P<0.0001*) and group x time interaction (F _(1, 26)_ = 20.79, *P<0.001*; Bonferroni post hoc (* = Saline vs ketamine; # = day 1 vs day 5), showing that ketamine induces hyperlocomotion. (e) Sub-chronic ketamine induces locomotor sensitization (\*\*\**P<0.001*). (f) Locomotor sensitization is sustained following a two-day washout of ketamine. Ketamine induced significantly higher locomotor activity in mice that had received sub-chronic ketamine as compared to mice that had received saline for 5 days (\*\**P<0.01*). Data represent mean ± S.E.M. \*\*\**P<0.001*, \*\**P<0.01*. Abbreviations: PET-positron emission tomography, CT-computed tomography

### Chemogenetics model

In *DAT∷Cre* mice AAVs were stereortaxically targeted to the ventral tegmental area (VTA: anteroposterior [AP], −3.15mm, mediolateral [ML] ±0.40mm, dorsoventral [DV] −4.30mm) and the substantia nigra pars compacta (SNc: AP, −3.15mm, ML ±1.50mm, DV −4.30mm) (Figure 2b). In *PV∷Cre* mice viruses were stereotaxically injected in pre-limbic cortex (PLc: AP +1.94, ML ±0.45, DV −2.20) and in the ventral subiculum (vSub) of the hippocampus (vSub: AP −3.20, ML ±2.80, DV −4.30) (Figure 4b). The needle was left in place for 3min post injection. Following injections, the wound was sutured (Mersilk, 3-1 Ethicon). Two weeks following the surgeries, clozapine N-oxide (CNO) (0.1mg/kg and 0.5mg/kg, i.p) or saline was administered 30-min before the injection of ketamine or saline (Figures 2a, 4a). See supplementary methods for further details.

**Figure 2:**
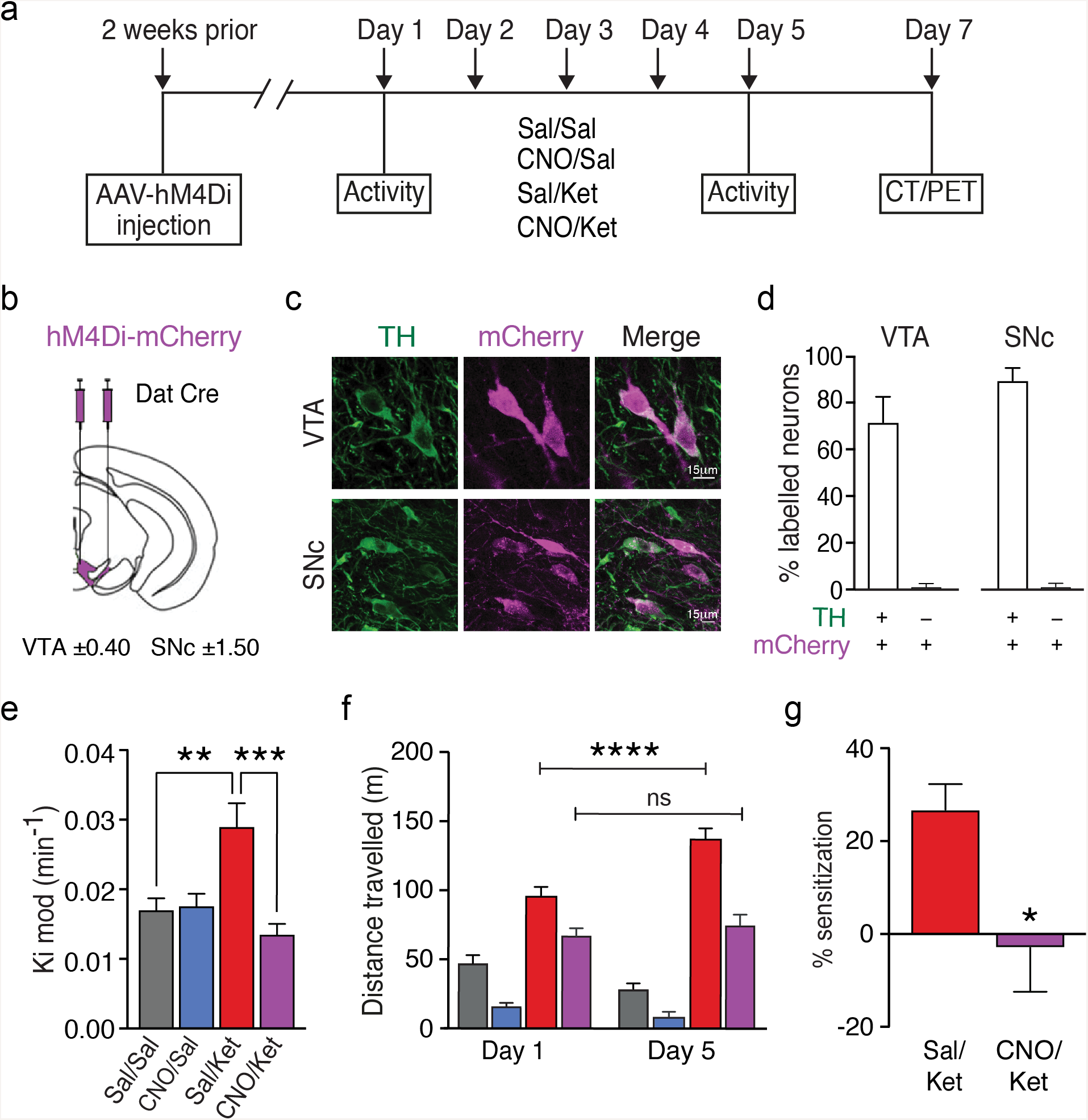
Midbrain dopamine neuron firing is necessary for ketamine-induced increases in dopamine synthesis capacity and locomotor activity. (a) Experimental timeline and drug treatment paradigm used to assess the effect of midbrain dopamine neuron inhibition on the sub-chronic ketamine-induced increase in striatal presynaptic dopamine synthesis capacity and locomotor activity. Two weeks after stereotaxic injection of AAV-hM4Di-mCherry, mice received 0.1mg/kg CNO or vehicle followed by ketamine (30mg/kg) or vehicle 30min later for 5 consecutive days. Mice underwent a dynamic PET/CT scan two days after the last drug administration. (b) Bilateral infusion of AAV-hM4Di-mCherry into the VTA and SNc of DAT-Cre mice was used to selectively express DREADD receptors in dopamine neurons. (c) Fluorescence confocal images of representative midbrain fields depicting coexpression (white) of mCherry (magenta) and TH immunofluorescence (green). (d) Percentage of TH+ neurons co-expressing mCherry (47 out of total 65 TH+ neurons; 71.7 ± 11 %) and percentage of mCherry+ which do not express TH (1 out of total 48 mCherry + neurons, 1.5 ± 1.5%) in the ventral tegmental area. Percentage of TH+ neurons co-expressing mCherry (47 out of total 52 TH+ neurons; 89.6% ± 5.4) and percentage of mCherry+ which do not express TH (1 out of total 48 mCherry+ neurons, 1.5 ± 1.5 %) in the SNc. (e) Striatal dopamine synthesis capacity (*K*_i_ ^mod^/minute) is significantly reduced in CNO/Ket compared to Sal/Ket group (\*\*\**P*<0.001) (Sal/Sal (*n*=12), CNO/Sal (*n*=13), Sal/Ket (*n*=12) and CNO/Ket-treated (*n*=11) groups). (f) Total distance travelled during 30min post drug administration. Sub-chronic ketamine treatment induced locomotor sensitization that was prevented by inhibition of midbrain dopamine neuron firing prior to ketamine treatment. (g) Percentage locomotor sensitization between day1 and day 5. Data represent mean ± S.E.M. \*\*\*\**P<0.0001*, \*\*\**P<0.001*, \*\**P<0.01*, \**P<0.05*. Abbreviations: Sal- saline, Ket- ketamine, CNO- clozapine N-oxide, TH- tyrosine hydroxylase, PET- positron emission tomography; CT- computed tomography; VTA- ventral tegmental area; SNc- substantia nigra pas compacta.

### Open-field test

Mice were placed into the open field arena for a 20-min habituation period, then injected i.p, with either ketamine or saline and placed back in the arena for a further 60-min. Total distance travelled was recorded using the Ethovision XT video tracking software system (Noldus Information Technologies, Leesburg, VA, USA). For the chemogenetic experiments, mice were placed in the open field arenas for 20-min habituation, they then received an injection of CNO or saline and activity was recorded for 30-min. Then, mice received an injection of ketamine or saline, in line with the treatment schedule, and their activity recorded for 60-min. Locomotor activity was assessed on days 1 and 5 of ketamine treatment.

### Positron emission tomography (PET) imaging

One hour prior to scanning, mice were anaesthetized with isoflurane and underwent external jugular vein cannulation. During scanning, the respiration rate was monitored using the BioVet physiological monitoring software system (Biovet software; m2m Imaging Corp, Cleveland, OH, USA) and body temperature was maintained at 37°C. Mice received 40mg/kg (i.p) entacapone (SML0654, Sigma-Aldrich), a catechol-O-methyl-transferase inhibitor, and 10mg/kg (i.p) benserazide hydrochloride (B7283, Sigma-Aldrich), an aromatic amino acid decarboxylase inhibitor, at 45-min and 30-min before the [^18^F]-FDOPA respectively. This improves brain uptake [^18^F]-FDOPA by reducing peripheral metabolism of the radiotracer (22). SEP363586 (3mg/kg, i.p) was provided by Sunovion Pharmaceuticals and administered 30-min prior to the [^18^F]-FDOPA injection. Following cannulation, mice were transferred to the bore of an Inveon μPET/CT scanner (Siemens, Surrey, UK). Mice underwent a 20-min CT scan for attenuation correction, and then received a bolus injection of approximately 4.5MBq [^18^F]-FDOPA via the external jugular vein cannula at the start of the 120-min dynamic PET scan.

### PET analysis

Inveon Research Workplace software (Siemens USA) was used to draw 3D regions of interest (ROIs) manually on summation radioactivity images at the level of the striatum (right and left) (0.07cm^3^) and the cerebellum (0.1cm^3^) to extract time activity curves (TACs) (Supplementary Figure 7) (23). Dopamine synthesis capacity was indexed as the rate constant for the uptake and conversion of [^18^F]-FDOPA to [^18^F]-dopamine, *K*_i_^mod^ (min^−1^), and determined using a modified Patlak plot accounting for the loss of radioactive metabolites, k_loss_ (22, 24). The cerebellum was used as the reference region, in line with the approach used in human studies, to account for non-specific uptake as it has negligible dopaminergic projections (25, 26).

### RNA Sequencing (RNA Seq)

Two days following 5 days of ketamine or saline injections, brains were rapidly removed and the PLc was dissected and frozen in isopentane on dry ice. Total RNA was isolated using the TriZol reagent (Invitrogen) and purified using RNAeasy micro kits from Qiagen. RNA integrity was assessed using the Agilent Bioanalyser and all RNA Integrity number (RIN) values were above 8. Then, the cDNA library was prepared using the NEB Next Ultra II Library Prep kit (New England Biolabs, USA). Sequencing was conducted on an Illumina HiSeq 2500 system with 100 base pair paired-end reads (London, UK). Raw reads were aligned to mm9 genome using Tophat version (2.0.11) (27). Gene based counting was performed using the HTSeq counts module. Gene expression analysis was performed using the DESeq2 Bioconductor package. All genes with adjusted p value of 0.05 or less (calculated from the raw p values using the Benjamini and Hochberg algorithm) were considered statistically significant.

### Statistical Analysis

Statistical analyses were performed using Prism 7.00 software (GraphPad Software, La Jolla, California, USA). Normality of distributions was assessed using the Kolmogorov-Smirnov test and Levene’s test for equality of variance to guide the choice of statistic. Between-group comparisons were made with two-tailed independent samples t-tests for normally distributed data, and Mann Whitney U tests were used for non-parametric data. Two-way analysis of variance (ANOVA) was used to test the difference in outcome measure between the four experimental groups. Locomotor sensitization was analysed with a two-way repeated-measures ANOVA, with the days as the repeated measure and experimental group as the cofactor. Outliers in the data were identified using the Grubbs’s test. Post hoc comparisons were Bonferroni corrected. Cohen’s d effect sizes were calculated using the online calculator (http://www.uccs.edu/~lbecker/). Data are expressed as mean ± s.e.m. and statistical significance was defined as p<0.05 (two-tailed).

## RESULTS

### Sub-chronic ketamine increases dopamine synthesis capacity and locomotor activity

To test the hypothesis that sub-chronic ketamine administration leads to increased dopamine synthesis capacity, mice were injected once daily with ketamine (30 mg/kg) or saline for 5 consecutive days. Two days after the last ketamine or saline injection *in vivo* [^18^F]-FDOPA PET imaging was performed. Sub-chronic ketamine treatment significantly increased striatal dopamine synthesis capacity compared to controls, with an effect size of d=2.5 (*P<0.001*, t_13_= 4.74) (Figure 1b, c, Supplementary Table 1). We also examined locomotor sensitization, which has been used as a behavioural model of the dopaminergic dysfunction seen in psychosis (28). Acute ketamine administration (day 1) induced locomotor hyperactivity in the open field test. Repeated ketamine administration (day 5) induced locomotor sensitization, an effect that was sustained following two-day washout of ketamine (day 7) (Figure 1d, e, f; Supplementary Figure 1). Collectively, these findings indicate that sub-chronic ketamine administration induces both an increase in dopamine synthesis capacity and behavioural changes relevant to schizophrenia.

### Midbrain dopamine neuron firing is necessary for ketamine-induced increases in dopamine synthesis capacity and locomotor activity

To test if the reported ketamine-induced firing activity of dopamine neurons (29–31), underlies the increase in dopamine synthesis capacity we observed, we employed a chemogenetic approach to selectively suppress dopamine neuron activity *in vivo*. We injected an adeno-associated virus (AAV) containing a Cre-dependent *hM4Di-mCherry* fusion protein (AAV1-DIO-hM4Di-mCherry) into the ventral tegmental area (VTA) and the substantia nigra pars compacta (SNc) of *DAT∷Cre* mice. Cre-dependent expression of *hM4Di-mCherry* showed ^~^98% specificity for dopamine neurons, and CNO-treatment silenced dopamine neuron firing in slice electrophysiology recordings, consistent with our previous findings (Figure 2b-d)(32). Administration of CNO prior to ketamine dosing prevented the elevation in striatal dopamine synthesis capacity (Figure 2e, Supplementary Table 2) and the ketamine-induced locomotor sensitization compared to the relevant control groups (Figure 2f&g, Supplementary Fig.2). It has recently been shown that clozapine, converted from CNO, may have off-target effects at endogenous receptors rather than at the DREADDs exclusively (33). Importantly, CNO administration in transgenic mice expressing a control construct had no effect on the ketamine-induced increase in dopamine synthesis capacity and locomotor activity (Supplementary Figure 4), indicating that DREADD-mediated silencing of dopaminergic neurons is responsible for the observed effects. Taken together, these findings suggest that sub-chronic ketamine increases dopamine synthesis capacity and locomotor sensitization through a mechanism that drives firing activity of midbrain dopamine neurons.

### The effect of sub-chronic ketamine on PV expression and function

Lower levels of PV neurons in the cortex and hippocampus have been observed in schizophrenia patients and following acute ketamine treatment (16, 17, 34, 35). In addition, it is believed that reduced PV neuron function may lead to changes in dopamine neuron activity (19). Therefore, we examined the effects of ketamine on various elements of PV interneuron function including PV expression. We found that sub-chronic ketamine treatment reduced PV interneuron immunofluorescence in the pre-limbic cortex (PLc) and the ventral subiculum (vSub) of the hippocampus (*P<0.05*, η^2^ effect size= 0.86) relative to saline controls (Figure 3a).

**Figure 3:**
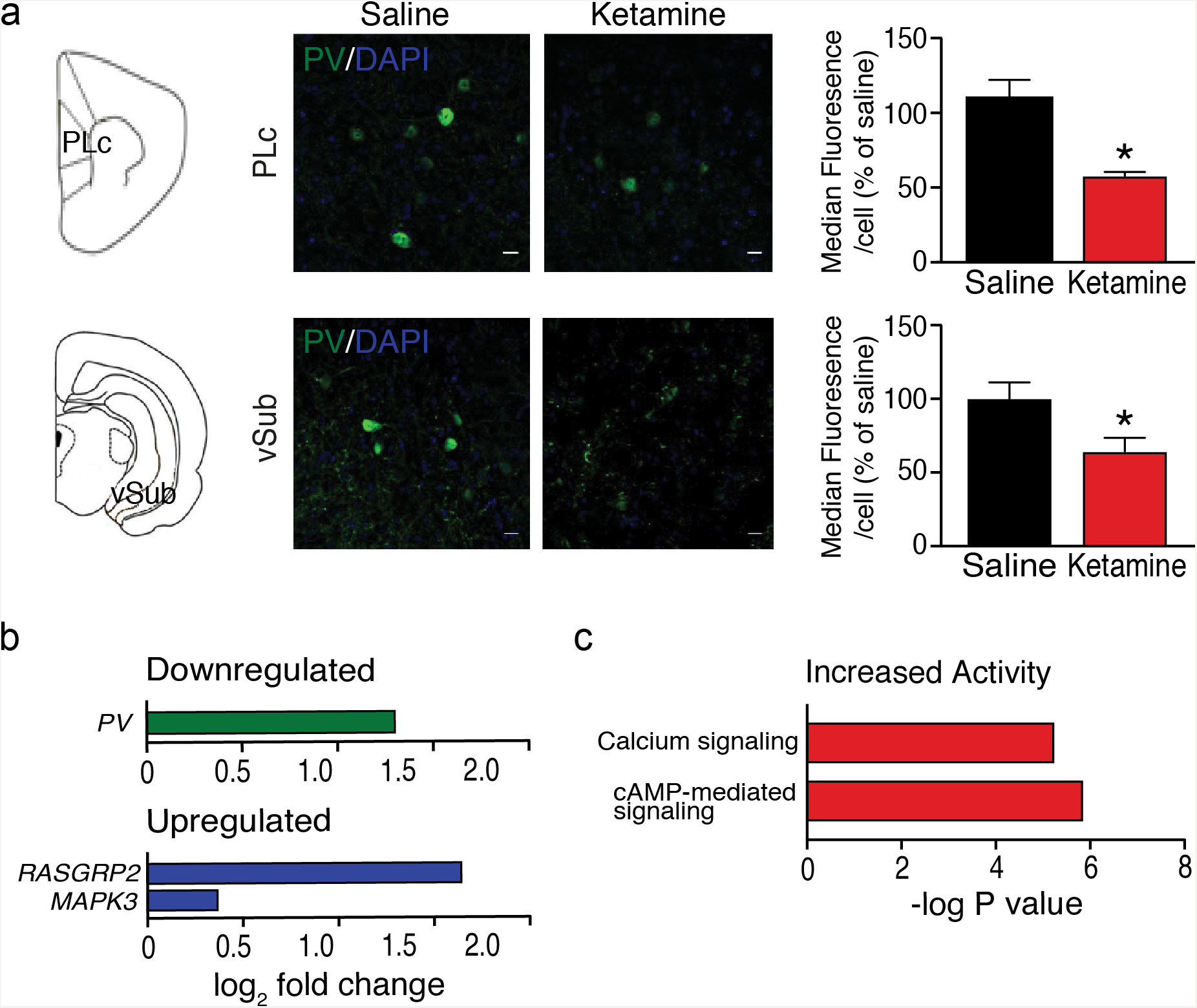
Sub-chronic ketamine reduces parvalbumin interneuron function. (a) Schematics of the location of the PLc and vSub of the hippocampus in the brain. Representative fluorescence confocal images of pre-limbic cortex and ventral subiculum (vSub) of the hippocampus fields respectively depicting PV interneuron immunofluorescence (green) in control (Control) and ketamine-treated (Ketamine) mice. PV immunofluorescence in the PLc and vSub of the hippocampus is significantly reduced in the ketamine versus saline group (Two-way repeated measures ANOVA significant effect of treatment, F_1, 8_= 47.28, p<0.001, η^2^ Effect size= 0.86; followed by Bonferroni post hoc tests (*P*<0.05); *n*=5 mice per group). (b) Differential expression of the PV gene in ketamine treated mice vs saline treated controls. Differential expression of RASGRP2 and MAPK3 genes in ketamine treated mice vs saline treated controls. Log2 fold change is shown in each respective bar. (c) Increased activity in the calcium signalling and cAMP-mediated signalling pathways in ketamine-treated vs control group. Data represent mean ± S.E.M. \**P<0.05*. Abbreviations: PLc-pre-limbic cortex; vSub-ventral subiculum of the hippocampus; PV-parvalbumin.

To investigate the molecular mechanisms underlying the effects of sub-chronic ketamine on dopamine synthesis we performed RNA sequencing (RNA Seq) on Plc tissue. We hypothesized that sub-chronic ketamine would result in reduced PV expression, and changes in signalling pathways downstream of the NMDA receptor such as calcium signalling and the activation of BDNF signalling. Consistent with our a-priori hypotheses, RNA Seq data on PLc tissue revealed reductions in the expression of PV (Figure 3b). Moreover, consistent with the hypothesis that blocking NMDAR activity increases BDNF signalling (36), we observed a significant increase in the expression of genes involved in the pathway downstream of BDNF signalling, specifically upregulation of mitogen-activated protein kinase 3 (MAPK3) and RAS, guanyl releasing protein 2 (Rasgrp2) (Figure 3b). Additionally, cAMP mediated signalling and calcium signalling pathways were significantly activated in ketamine vs saline conditions (Figure 3c). Furthermore, using Ingenuity pathway analysis (IPA, QUIAGEN Redwood City, https://www.qiagenbioinformatics.com/products/ingenuity-pathway-analysis/) L-DOPA was the significant upstream regulator of the differentially expressed genes in ketamine vs saline conditions (z-score= 2.961, *P<0.05*, in cortex). Collectively, these data support the hypothesis that sub-chronic ketamine increases dopamine synthesis capacity via a pathway that involves the inhibition of PV interneuron function.

### The role of PV interneuron activity in mediating the effects of sub-chronic ketamine

Given that ketamine reduced PV expression levels, and the hypothesised role of PV interneuron hypofunction in schizophrenia (37, 38), we aimed to determine if activating cortical PV interneurons was able to counter the ketamine-induced increase in dopamine synthesis capacity. To test this, AAVs expressing a Cre-dependent *hM3Dq-mCherry* fusion protein were stereotaxically injected into the PLc and vSub of the hippocampus of *PV∷Cre* mice (Figure 4a&b). Immunohistochemistry revealed co-localization of mCherry with PV immunoreactive neurons and a successful transduction with over 92% specificity in the PLc and vSub (Figure 4c&d). In *ex vivo* slice electrophysiology studies, application of CNO depolarised vSub PV neurons expressing mCherry (Figure 4e&f). Using this system, we found that *in vivo* activation of PV interneurons in the PLc and vSub, prior to ketamine administration, significantly reduced both the elevation in striatal dopamine synthesis capacity (*P*<0.01, t_19_= 3.51, two-tailed, Effect size= 1.59) (Figure 4g; Supplementary Table 3) and the locomotor effects of acute and sub-chronic ketamine (Figure 4h; Supplementary Figure 3). Our results, therefore, suggest that ketamine increases dopamine synthesis capacity and locomotor activity via its actions on cortical/hippocampal PV interneurons.

**Figure 4:**
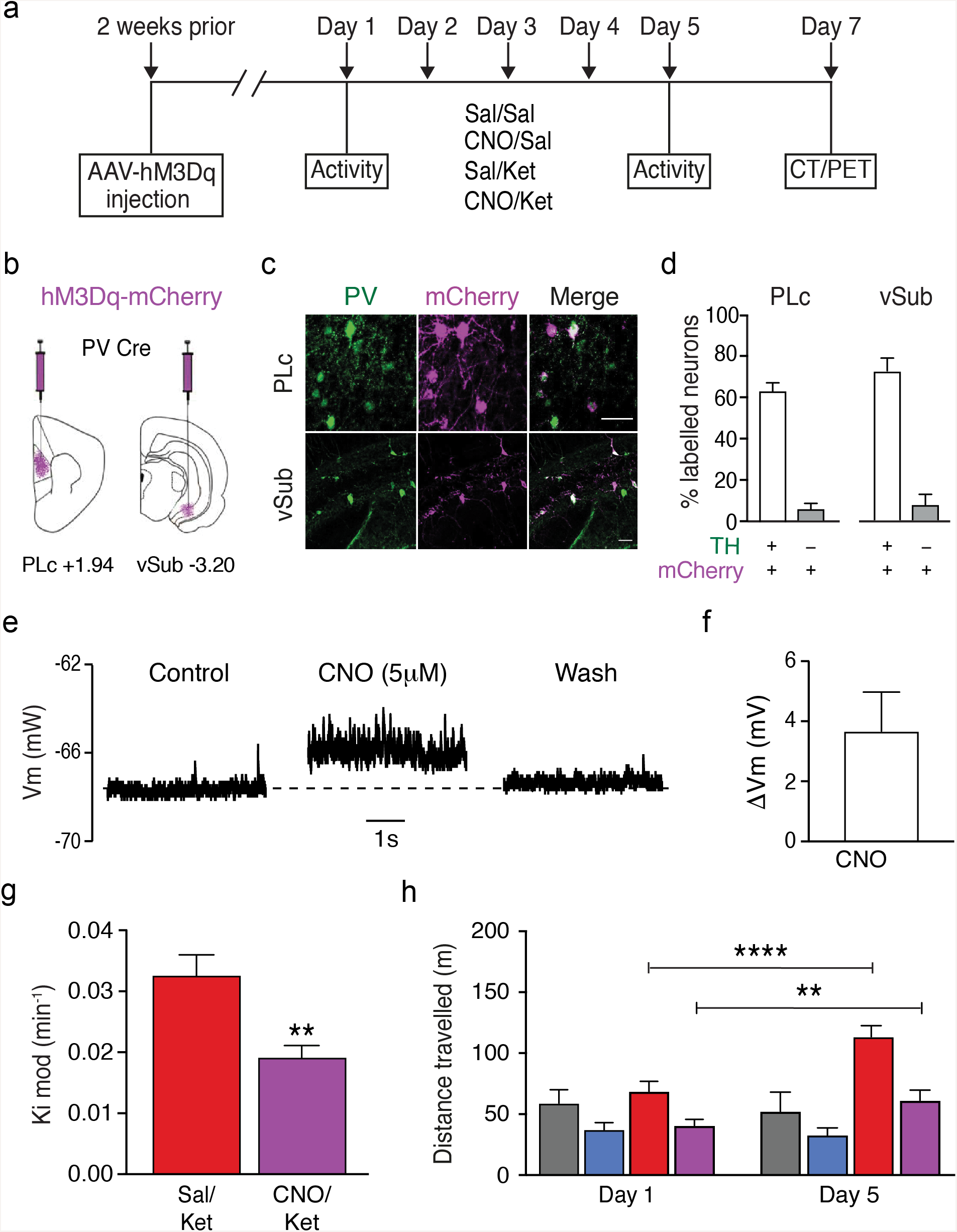
*in vivo* parvalbumin interneuron activation attenuates the effects of sub-chronic ketamine-induced increase in dopamine synthesis capacity and locomotor activity. (a) Experimental timeline and drug treatment schedule used to study the effect of PV neuron activation on the sub-chronic ketamine-induced increase in striatal presynaptic dopamine synthesis capacity. Two weeks following stereotaxic injection of AAV-hM3Dq-mCherry, mice received 0.5mg/kg CNO or vehicle, followed by ketamine (30mg/kg) or vehicle treatment 30min later for 5 consecutive days. Mice underwent a dynamic PET/CT scan two days following the last drug administration. (b) Bilateral infusion of AAV-hM3Dq-mCherry into the PLc and vSub of PV-Cre mice was used to selectively express DREADD receptors in parvalbumin interneurons. (c) Representative fluorescence confocal images of PLc and vSub fields depicting coexpression (white) of mCherry (magenta) and parvalbumin immunofluorescence (green). (d) Percentage of PV+ neurons co-expressing mCherry (120 out of total 177 PV+ neurons; 65 ± 4.4%) and percentage of mCherry+ which do not express PV (9 out of total 129 mCherry+ neurons, 5.8 ± 2.9 %) in the PLc. Percentage of PV+ neurons co-expressing mCherry (51 out of total 71 PV+ neurons; 72.9 ± 6.3 %) and percentage of mCherry+ which do not express PV (5 out of total 59 mCherry + neurons, 7.8 ± 5.2%) in the vSub. (e) Effect of 5mM CNO on membrane potential measured in voltage clamp configuration from a whole-cell recording of PV interneuron within the vSub from a PV-Cre mouse injected with AAV-hM3Dq-mCherry. (f) Change in membrane potential with positive showing increase relative to the baseline indicative of PV neuron depolarization upon CNO application. (g) Striatal dopamine synthesis capacity (*K*_i_ ^*mod*^/minute) is significantly reduced in CNO/ketamine-treated (*n=11*) (purple) versus SAL/Ket (*n=11*) (red) group, unpaired t-test (\*\**P<0.01*, t_19_= 3.51, two-tailed, Effect size= 1.59). (h) Total distance travelled during a 30min test period post drug administration. Sub-chronic ketamine treatment induced locomotor sensitization, which was not prevented by activation of parvalbumin interneuron firing prior to ketamine treatment (F_1, 44_ =15.51, *** *P<0.001*, Bonferroni post-hoc test ** *P<0.01* *** *P<0.001* = day 1 vs day 5). On day 5 activation of parvalbumin interneuron firing prevented the effects of sub-chronic ketamine on locomotor activity (F _3, 44_ = 9.283, \*\*\**P<0.001*, Bonferroni post-hoc test \*\*\**P<0.001* Sal/Ket vs all other groups). Data represent mean ± S.E.M. \*\*\*\**P<0.001*, \*\**P<0.01*. Abbreviations: Sal- saline, Ket- ketamine, CNO- clozapine N-oxide, PET- positron emission tomography; CT- computed tomography; PLc- pre-limbic cortex, vSub- ventral subiculum of the hippocampus

### A novel TAAR1 agonist counteracts the ketamine-induced increase in dopamine synthesis capacity

Our findings suggest that targeting dopamine neuron firing activity may present a viable therapeutic target for the increase in dopamine synthesis capacity seen in schizophrenia. One potential candidate mechanism is TAAR1 agonism. TAAR1 is a G protein-coupled receptor that is expressed throughout in monoaminergic brain nuclei including dopamine neurons (29). TAAR1 agonists have been shown to reduce dopamine firing rates and release (30–32). In view of this, we tested whether SEP-0363856 (SEP856), a novel psychotropic agent with agonism at TAAR1 (20), counteracts the effect of sub-chronic ketamine treatment on dopamine synthesis capacity. Ketamine-treated mice that received SEP856 (3mg/kg, i.p) showed significantly lower striatal dopamine synthesis capacity compared to vehicle controls (P<0.01, t36= 3.41) (Figure 5). Post hoc analyses showed that Ki mod in ketamine treated mice that received SEP856 was not significantly different from Ki mod in mice not exposed to ketamine (Figure 5).

**Figure 5:**
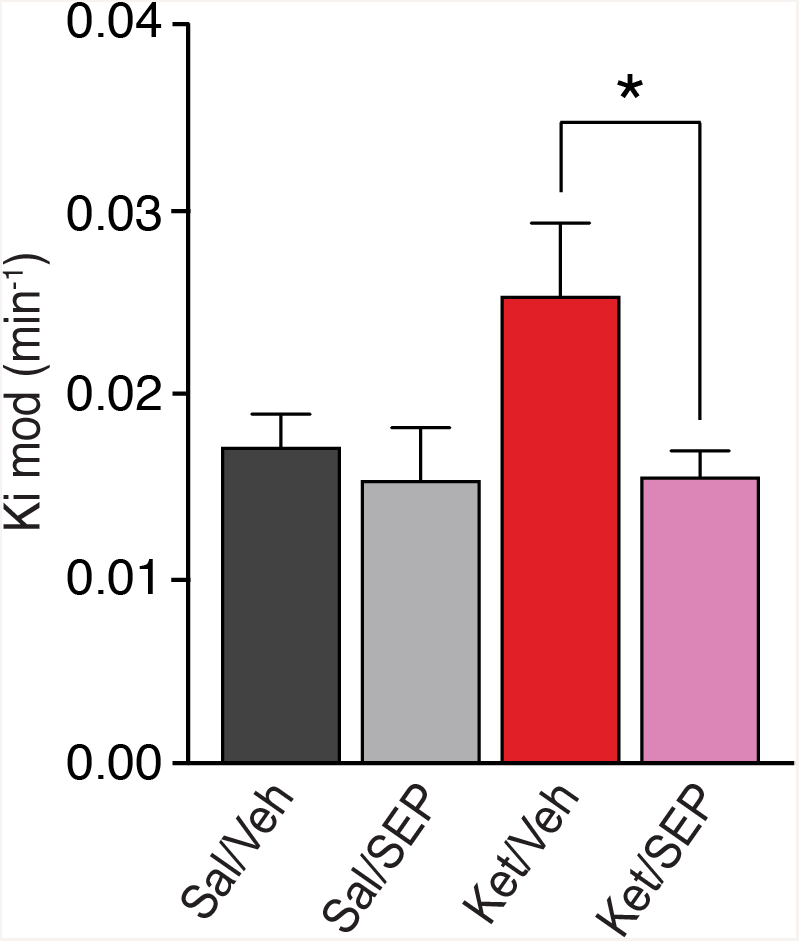
The novel trace amine receptor 1 agonist SEP-0363856 (SEP856) attenuates the ketamine-induced increase in dopamine synthesis capacity. Striatal dopamine synthesis capacity (*K*_i_ ^*mod*^/minute) was significantly reduced in the ketamine model following the administration of SEP compared to vehicle in the ketamine model. Ket/SEP (*n=9*) (Magenta) versus Ket/Veh (*n=8*) (red) group \**P<0.05*. Abbreviations: Sal- saline, Ket- ketamine, Veh- vehicle, SEP- SEP-0363856

## DISCUSSION

Our results demonstrate that sub-chronic ketamine administration leads to elevated striatal dopamine synthesis capacity, which requires the activation of midbrain dopamine neurons. The ketamine-induced increase in dopamine synthesis capacity was attenuated by the activation of parvalbumin interneurons in the pre-limbic cortex and ventral subiculum, as well as by a novel TAAR1 agonist, SEP856.

The majority of studies show that acute ketamine significantly increases striatal dopamine levels (39–42), and elevates VTA dopamine neuron firing (30, 31, 43). Our results extend these findings to show that sub-chronic ketamine induces a persistent elevation of dopamine synthesis capacity through a mechanism that requires midbrain dopamine neuron firing. These results are in line with the increased striatal dopamine synthesis capacity observed in schizophrenia patients and were acquired using an equivalent PET imaging technique. Additionally, we extend prior findings of reductions in parvalbumin levels in the hippocampus and prefrontal cortex following acute or sub-chronic ketamine administration (34, 35), to show that ketamine also leads to persistent reductions in PV levels and that activation of PV interneurons can attenuate ketamine-induced increases in striatal dopamine synthesis capacity.

### Proposed mechanism of action of ketamine on dopamine synthesis capacity

Ketamine is a non-competitive NMDAR antagonist that binds with high affinity Ki= 3.1 μM to the same binding site as MK801 and PCP (44–46). Parvalbumin interneurons are activated upon glutamate binding to NMDAR, and subsequently inhibit the activity of cortical pyramidal neurons (47–49). Therefore, by blocking NMDAR, ketamine is thought to reduce the activity of parvalbumin interneurons and thereby disinhibit cortical pyramidal neurons, including neurons that project to subcortical regions to ultimately disinhibit midbrain dopamine neuron firing (19, 30, 50–52). In line with this model of the mechanism of action of ketamine on the dopaminergic system, ketamine has been associated with reduction in parvalbumin interneuron function, excessive glutamate release (53–56) and an increase in dopamine neuron firing (30, 31, 43). Our findings that ketamine’s effects can be reduced by activating parvalbumin interneurons and inhibiting midbrain dopamine neurons is consistent with this model. However, it remains possible that ketamine has actions on other neuronal populations that contribute to its effects on striatal dopamine synthesis capacity. Further work is required to test whether pyramidal neuron activity is altered in our ketamine model, and to characterise the circuit linking cortical parvalbumin interneurons to midbrain dopamine neurons. Ketamine’s action on receptors other than the NMDAR, could also contribute to the observed effects. Recent findings indicate that activation of dopamine D1 receptors on pyramidal cells in the prefrontal cortex and/or the action of a metabolite of ketamine on α-amino-3-hydroxy-5-methyl-4-isoxazolepropionic acid (AMPA) receptors might contribute to its long-lasting antidepressant effects (57) (58). However, ketamine’s affinities at other receptors (range of Ki values= 19-131μM) are considerably lower than its affinity for the NMDAR, and it is not clear if ketamine exhibits significant striatal dopamine receptor occupancy *in vivo* at behaviourally relevant doses (59, 60). We suggest that the effects of ketamine in our model likely involve NMDAR blockade, but a contribution from binding to other receptors cannot be excluded.

Ketamine has a short half-life (13min) in mice (61) and brain levels of its main metabolite (2R, 6R)-hydroxynorketamine (HNK), are not detectable 4-hours post administration in mice (58, 62, 63). Our PET and behavioural measures were acquired ^~^48 hours following the last ketamine treatment. Thus, it is unlikely that the observed effects are a consequence of direct ketamine or HNK action. Previous studies have shown that ketamine induces a release of brain derived neurotrophic factor (BDNF) to increase synaptogenesis (63) and elevates MAPK signalling (64, 65). This could present a potential mechanism by which ketamine contributes to the sustained effects observed in our model.

A strength of our study is that it utilized a PET imaging approach that parallels the technique used in human studies (6, 22, 25), supporting the translational relevance of the findings. One consideration for chemogenetic approaches is the cell-type and regional specificity of expression. Injection of viral constructs in wild type mice revealed no detectable expression, and CNO administration in transgenic mice expressing a control construct had no effect on the ketamine-induced increase in dopamine synthesis capacity and locomotor activity. Limitations include that we did not measure other aspects of dopamine function or investigate other brain regions. Moreover, we did not test whether the effects of SEP-856 are predominantly mediated via TAAR1 agonism and its action on dopamine neuron firing, or potentially also driven by the compound’s activities at other receptors including 5-HT1A (20). It will be useful to integrate these considerations into future studies.

### Implications for understanding the pathophysiology of schizophrenia and the antidepressant mode of action of ketamine

PET imaging studies have repeatedly shown elevated dopamine synthesis and release capacity in schizophrenia (e.g. (7, 66) and see review (6)), and suggested that this is linked to the development of psychosis (67, 68) and changes in cortical glutamate levels (69). Moreover, ketamine induces psychotic symptoms in healthy volunteers and worsens symptoms in patients with schizophrenia (14, 15). Our findings reveal the underlying circuit by which ketamine induces elevated dopamine synthesis capacity, which is relevant to our understanding of the clinical observations. These findings suggest that inhibition of midbrain dopamine neurons and/or activation of cortical parvalbumin interneurons could represent novel therapeutic strategies for schizophrenia. Furthermore, our finding that SEP856, a novel TAAR1 agonist, reduces sub-chronic ketamine-induced elevation in striatal dopamine synthesis capacity provides a proof-of-concept for pharmacological attenuation of presynaptic dopamine dysfunction.

Lastly, our data also have implications for understanding ketamine’s antidepressant actions and its abuse potential. There is some evidence that major depression is associated with blunted dopaminergic function, including reduced levels of dopamine metabolites post-mortem and reduced dopamine synthesis capacity (70, 71). Moreover, animal models mimicking the neurochemical changes seen in depression exhibit reduced dopamine neuron population activity (30). Our findings that sub-chronic ketamine administration elevates striatal dopamine synthesis capacity, which persists for several days post dosing, suggest that this could contribute to ketamine’s antidepressant actions (48). Furthermore, a significant reduction in cortical ERK1/2 MAPK activity, protein and mRNA level were reported in major depressive disorder (MDD)-suicide patients (72). In line with previous findings, we show that sub-chronic ketamine administration leads to the elevation of MAPK signalling, suggesting this could contribute to ketamine’s antidepressant effects (64, 65).

## Conclusion

We demonstrate that sub-chronic ketamine leads to an increase in striatal dopamine synthesis capacity in the mouse, resembling the dopaminergic alteration seen in patients with schizophrenia. Our data suggest that ketamine’s effects on dopamine synthesis capacity are mediated by inhibition of PV interneurons in the cortex and ventral subiculum as well as activation of midbrain dopamine neurons, and that these alterations can be attenuated by agonism of TAAR1. Together, our findings provide novel insights into neurocircuits that drive dopaminergic dysfunction thought to underlie psychosis, which is of relevance for the development of new therapeutic approaches.

## Acknowledgments

We are grateful to Darran Hardy, Paulius Viskaitis, Maria Paiva-Pessoa, Sac-Pham Tang, Sharon Ashworth, Bilbinder Bhachu, Anna Pacelli, Stuart McCluskey, and the radiochemistry and biology teams at Invicro and to the CBS staff members at Imperial College London and the genomics facility and bioinformatics facility at the London Institute of Medical Sciences. The dopamine transporter *(DAT)∷Cre* mouse line was kindly gifted by Professor Francois Tronche (University of Pierre and Marie Curie, Paris). The study was funded by the Medical Research Council (MRC) UK grant to Professor Howes (MC-A656-5QD30), Professor Withers (MC-A654-5QB40) and Dr Ungless (MC-A654-5QB70).

## Author contributions

M.K, O.D.H, M.A.U, D.J.W and E.E.I contributed to study design. M.K, E.E.I, D.R.B, S.N, L.A.W, M.A.S, J.G., E.J.P, K.T., M.V, S.K, N.D, S.C.H, M.A.U, D.J.W and O.D.H. contributed to data collection or interpretation. M.K coordinated all experiments. M.K., D.J.W and O.D.H. wrote the original draft. All the authors critically revised the article. All authors approved the last version. Supervision was provided by M.A.U, D.J.W and O.D.H.

## References

1. Whiteford HA, Degenhardt L, Rehm J, Baxter AJ, Ferrari AJ, Erskine HE, et al. Global burden of disease attributable to mental and substance use disorders: findings from the Global Burden of Disease Study 2010. Lancet (London, England). 2013;382(9904):1575–86.

2. Howes O, McCutcheon R, Stone J. Glutamate and dopamine in schizophrenia: an update for the 21st century. Journal of psychopharmacology. 2015;29(2):97–115.

3. Lodge DJ, Grace AA. Hippocampal dysregulation of dopamine system function and the pathophysiology of schizophrenia. Trends in pharmacological sciences. 2011;32(9):507–13.

4. Weinstein JJ, Chohan MO, Slifstein M, Kegeles LS, Moore H, Abi-Dargham A. Pathway-Specific Dopamine Abnormalities in Schizophrenia. Biological psychiatry. 2017;81(1):31–42.

5. Uchida H, Takeuchi H, Graff-Guerrero A, Suzuki T, Watanabe K, Mamo DC. Dopamine D2 receptor occupancy and clinical effects: a systematic review and pooled analysis. Journal of clinical psychopharmacology. 2011;31(4):497–502.

6. Howes OD, Kambeitz J, Kim E, Stahl D, Slifstein M, Abi-Dargham A, et al. The nature of dopamine dysfunction in schizophrenia and what this means for treatment. Archives of general psychiatry. 2012;69(8):776–86.

7. Reith J, Benkelfat C, Sherwin A, Yasuhara Y, Kuwabara H, Andermann F, et al. Elevated dopa decarboxylase activity in living brain of patients with psychosis. Proceedings of the National Academy of Sciences of the United States of America. 1994;91(24):11651–4.

8. Hietala J, Syvalahti E, Vuorio K, Rakkolainen V, Bergman J, Haaparanta M, et al. Presynaptic dopamine function in striatum of neuroleptic-naive schizophrenic patients. Lancet (London, England). 1995;346(8983):1130–1.

9. Meyer-Lindenberg A, Miletich RS, Kohn PD, Esposito G, Carson RE, Quarantelli M, et al. Reduced prefrontal activity predicts exaggerated striatal dopaminergic function in schizophrenia. Nature neuroscience. 2002;5(3):267–71.

10. Howes O, Bose S, Turkheimer F, Valli I, Egerton A, Stahl D, et al. Progressive increase in striatal dopamine synthesis capacity as patients develop psychosis: a PET study. Molecular psychiatry. 2011;16(9):885–6.

11. Jauhar S, Nour MM, Veronese M, Rogdaki M, Bonoldi I, Azis M, et al. A Test of the Transdiagnostic Dopamine Hypothesis of Psychosis Using Positron Emission Tomographic Imaging in Bipolar Affective Disorder and Schizophrenia. Jama Psychiat. 2017;74(12):1206–13.

12. Abi-Dargham A, Laruelle M. Mechanisms of action of second generation antipsychotic drugs in schizophrenia: insights from brain imaging studies. European psychiatry: the journal of the Association of European Psychiatrists. 2005;20(1):15–27.

13. Howes OD, McCutcheon R, Owen MJ, Murray RM. The Role of Genes, Stress, and Dopamine in the Development of Schizophrenia. Biological psychiatry. 2017;81(1):9–20.

14. Krystal JH, Karper LP, Seibyl JP, Freeman GK, Delaney R, Bremner JD, et al. Subanesthetic effects of the noncompetitive NMDA antagonist, ketamine, in humans. Psychotomimetic, perceptual, cognitive, and neuroendocrine responses. Archives of general psychiatry. 1994;51(3):199–214.

15. Lahti AC, Koffel B, LaPorte D, Tamminga CA. Subanesthetic doses of ketamine stimulate psychosis in schizophrenia. 1995;13(1):9–19.

16. Benes FM, McSparren J, Bird ED, SanGiovanni JP, Vincent SL. Deficits in small interneurons in prefrontal and cingulate cortices of schizophrenic and schizoaffective patients. Archives of general psychiatry. 1991;48(11):996–1001.

17. Zhang ZJ, Reynolds GP. A selective decrease in the relative density of parvalbumin-immunoreactive neurons in the hippocampus in schizophrenia. Schizophr Res. 2002;55(1-2):1–10.

18. Dwir D, Giangreco B, Xin L, Tenenbaum L, Cabungcal JH, Steullet P, et al. MMP9/RAGE pathway overactivation mediates redox dysregulation and neuroinflammation, leading to inhibitory/excitatory imbalance: a reverse translation study in schizophrenia patients. Molecular psychiatry. 2019;Epub ahead of print.

19. Grace AA. Dysregulation of the dopamine system in the pathophysiology of schizophrenia and depression. Nature reviews Neuroscience. 2016;17(8):524–32.

20. Dedic N, Jones PG, Hopkins SC, Lew R, Shao L, Campbell JE, et al. SEP-363856, A novel psychotropic agent with a unique, non-D2 receptor mechanism of action. The Journal of pharmacology and experimental therapeutics. 2019.

21. McNally JM, McCarley RW, Brown RE. Chronic Ketamine Reduces the Peak Frequency of Gamma Oscillations in Mouse Prefrontal Cortex Ex vivo. Frontiers in psychiatry. 2013;4:106.

22. Walker MD, Dinelle K, Kornelsen R, McCormick S, Mah C, Holden JE, et al. In-vivo measurement of LDOPA uptake, dopamine reserve and turnover in the rat brain using [18F]FDOPA PET. Journal of cerebral blood flow and metabolism: official journal of the International Society of Cerebral Blood Flow and Metabolism. 2013;33(1):59–66.

23. Bonsall DR, Kokkinou M, Veronese M, Coello C, Wells LA, Howes OD. Single cocaine exposure does not alter striatal presynaptic dopamine function in mice: an [18 F]-FDOPA PET study. Journal of neurochemistry. 2017.

24. Holden JE, Doudet D, Endres CJ, Chan GL, Morrison KS, Vingerhoets FJ, et al. Graphical analysis of 6-fluoro-L-dopa trapping: effect of inhibition of catechol-O-methyltransferase. Journal of nuclear medicine: official publication, Society of Nuclear Medicine. 1997;38(10):1568–74.

25. Cumming P, Kuwabara H, Ase A, Gjedde A. Regulation of DOPA decarboxylase activity in brain of living rat. Journal of neurochemistry. 1995;65(3):1381–90.

26. Egerton A, Demjaha A, McGuire P, Mehta MA, Howes OD. The test-retest reliability of 18F-DOPA PET in assessing striatal and extrastriatal presynaptic dopaminergic function. Neuroimage. 2010;50(2):524–31.

27. Kim D, Pertea G, Trapnell C, Pimentel H, Kelley R, Salzberg SL. TopHat2: accurate alignment of transcriptomes in the presence of insertions, deletions and gene fusions. Genome biology. 2013;14(4):R36.

28. O'Tuathaigh CM, Waddington JL. Closing the translational gap between mutant mouse models and the clinical reality of psychotic illness. Neurosci Biobehav Rev. 2015;58:19–35.

29. French ED, Ceci A. Non-competitive N-methyl-D-aspartate antagonists are potent activators of ventral tegmental A10 dopamine neurons. Neuroscience letters. 1990;119(2):159–62.

30. Belujon P, Grace AA. Restoring mood balance in depression: ketamine reverses deficit in dopamine-dependent synaptic plasticity. Biological psychiatry. 2014;76(12):927–36.

31. Witkin JM, Monn JA, Schoepp DD, Li X, Overshiner C, Mitchell SN, et al. The Rapidly Acting Antidepressant Ketamine and the mGlu2/3 Receptor Antagonist LY341495 Rapidly Engage Dopaminergic Mood Circuits. The Journal of pharmacology and experimental therapeutics. 2016;358(1):71–82.

32. Sandhu EC, Fernando ABP, Irvine EE, Tossell K, Kokkinou M, Glegola J, et al. Phasic Stimulation of Midbrain Dopamine Neuron Activity Reduces Salt Consumption. eNeuro. 2018;5(2).

33. Gomez JL, Bonaventura J, Lesniak W, Mathews WB, Sysa-Shah P, Rodriguez LA, et al. Chemogenetics revealed: DREADD occupancy and activation via converted clozapine. Science. 2017;357(6350):503–7.

34. Zhou Z, Zhang G, Li X, Liu X, Wang N, Qiu L, et al. Loss of phenotype of parvalbumin interneurons in rat prefrontal cortex is involved in antidepressant-and propsychotic-like behaviors following acute and repeated ketamine administration. Molecular neurobiology. 2015;51(2):808–19.

35. Keilhoff G, Becker A, Grecksch G, Wolf G, Bernstein HG. Repeated application of ketamine to rats induces changes in the hippocampal expression of parvalbumin, neuronal nitric oxide synthase and cFOS similar to those found in human Schizophrenia. Neuroscience. 2004;126(3):591–8.

36. Gideons ES, Kavalali ET, Monteggia LM. Mechanisms underlying differential effectiveness of memantine and ketamine in rapid antidepressant responses. Proceedings of the National Academy of Sciences of the United States of America. 2014;111(23):8649–54.

37. Edwards AC, Bacanu SA, Bigdeli TB, Moscati A, Kendler KS. Evaluating the dopamine hypothesis of schizophrenia in a large-scale genome-wide association study. Schizophr Res. 2016;176(2-3):136–40.

38. Chung DW, Fish KN, Lewis DA. Pathological Basis for Deficient Excitatory Drive to Cortical Parvalbumin Interneurons in Schizophrenia. The American journal of psychiatry. 2016;173(11):1131–9.

39. Kokkinou M, Ashok AH, Howes OD. The effects of ketamine on dopaminergic function: meta-analysis and review of the implications for neuropsychiatric disorders. Molecular psychiatry. 2018;23(1):59–69.

40. Verma A, Moghaddam B. NMDA receptor antagonists impair prefrontal cortex function as assessed via spatial delayed alternation performance in rats: Modulation by dopamine. Jan 1996. The Journal of Neuroscience. 1996;.16(1):pp.

41. Usun Y, Eybrard S, Meyer F, Louilot A. Ketamine increases striatal dopamine release and hyperlocomotion in adult rats after postnatal functional blockade of the prefrontal cortex. Behavioural brain research. 2013;256:229–37.

42. Lai CC, Lee LJ, Yin HS. Combinational effects of ketamine and amphetamine on behaviors and neurotransmitter systems of mice. Neurotoxicology. 2013;37:136–43.

43. El Iskandrani KS, Oosterhof CA, El Mansari M, Blier P. Impact of subanesthetic doses of ketamine on AMPA-mediated responses in rats: An in vivo electrophysiological study on monoaminergic and glutamatergic neurons. Journal of Psychopharmacology. 2015;.29(7):pp.

44. Nishimura M, Sato K, Okada T, Yoshiya I, Schloss P, Shimada S, et al. Ketamine inhibits monoamine transporters expressed in human embryonic kidney 293 cells. Anesthesiology. 1998;88(3):768–74.

45. Seeman P, Ko F, Tallerico T. Dopamine receptor contribution to the action of PCP, LSD and ketamine psychotomimetics. Molecular psychiatry. 2005;10(9):877–83.

46. Dingledine R, Borges K, Bowie D, Traynelis SF. The glutamate receptor ion channels. Pharmacological reviews. 1999;51(1):7–61.

47. Javitt DC. Glutamatergic theories of schizophrenia. The Israel journal of psychiatry and related sciences. 2010;47(1):4–16.

48. Coyle JT. NMDA receptor and schizophrenia: a brief history. Schizophrenia bulletin. 2012;38(5):920–6.

49. Coyle JT. Glutamate and schizophrenia: beyond the dopamine hypothesis. Cellular and molecular neurobiology. 2006;26(4-6):365–84.

50. Javitt DC. Glutamate and schizophrenia: phencyclidine, N-methyl-D-aspartate receptors, and dopamine-glutamate interactions. International review of neurobiology. 2007;78:69–108.

51. Javitt DC, Zukin SR, Heresco-Levy U, Umbricht D. Has an angel shown the way? Etiological and therapeutic implications of the PCP/NMDA model of schizophrenia. Schizophrenia Bulletin. 2012;.38(5):pp.

52. Moghaddam B, Adams B, Verma A, Daly D. Activation of glutamatergic neurotransmission by ketamine: a novel step in the pathway from NMDA receptor blockade to dopaminergic and cognitive disruptions associated with the prefrontal cortex. The Journal of neuroscience: the official journal of the Society for Neuroscience. 1997;17(8):2921–7.

53. Rowland LM, Bustillo JR, Mullins PG, Jung RE, Lenroot R, Landgraf E, et al. Effects of ketamine on anterior cingulate glutamate metabolism in healthy humans: a 4-T proton MRS study. The American journal of psychiatry. 2005;162(2):394–6.

54. Stone JM, Dietrich C, Edden R, Mehta MA, De Simoni S, Reed LJ, et al. Ketamine effects on brain GABA and glutamate levels with 1H-MRS: relationship to ketamine-induced psychopathology. Molecular psychiatry. 2012;17(7):664–5.

55. Kim SY, Lee H, Kim HJ, Bang E, Lee SH, Lee DW, et al. in vivo and ex vivo evidence for ketamine-induced hyperglutamatergic activity in the cerebral cortex of the rat: Potential relevance to schizophrenia. Nmr Biomed. 2011;24(10):1235–42.

56. Abdallah CG, Sanacora G, Duman RS, Krystal JH. Ketamine and rapid-acting antidepressants: a window into a new neurobiology for mood disorder therapeutics. Annual review of medicine. 2015;66:509–23.

57. Hare BD, Shinohara R, Liu RJ, Pothula S, DiLeone RJ, Duman RS. Optogenetic stimulation of medial prefrontal cortex Drd1 neurons produces rapid and long-lasting antidepressant effects. Nature communications. 2019;10(1):223.

58. Zanos P, Moaddel R, Morris PJ, Georgiou P, Fischell J, Elmer GI, et al. NMDAR inhibition-independent antidepressant actions of ketamine metabolites. Nature. 2016;533(7604):481–6.

59. Can A, Zanos P. Effects of Ketamine and Ketamine Metabolites on Evoked Striatal Dopamine Release, Dopamine Receptors, and Monoamine Transporters. 2016;359(1):159–70.

60. Aalto S, Hirvonen J, Kajander J, Scheinin H, Nagren K, Vilkman H, et al. Ketamine does not decrease striatal dopamine D2 receptor binding in man. Psychopharmacology. 2002;164(4):401–6.

61. Maxwell CR, Ehrlichman RS, Liang Y, Trief D, Kanes SJ, Karp J, et al. Ketamine produces lasting disruptions in encoding of sensory stimuli. The Journal of pharmacology and experimental therapeutics. 2006;316(1):315–24.

62. Aleksandrova LR, Wang YT, Phillips AG. Hydroxynorketamine: Implications for the NMDA Receptor Hypothesis of Ketamine’s Antidepressant Action. Chronic stress (Thousand Oaks, Calif). 2017;1.

63. Abdallah CG. What’s the Buzz About Hydroxynorketamine? Is It the History, the Story, the Debate, or the Promise? Biological psychiatry. 2017;81(8):e61–e3.

64. Talebian A, Robinson-Brookes K, Meakin SO. TrkB Regulates N-Methyl-D-Aspartate Receptor Signaling by Uncoupling and Recruiting the Brain-Specific Guanine Nucleotide Exchange Factor, RasGrf1. Journal of molecular neuroscience: MN. 2019;67(1):97–110.

65. Reus GZ, Vieira FG, Abelaira HM, Michels M, Tomaz DB, dos Santos MA, et al. MAPK signaling correlates with the antidepressant effects of ketamine. Journal of psychiatric research. 2014;55:15–21.

66. Kumakura Y, Cumming P, Vernaleken I, Buchholz HG, Siessmeier T, Heinz A, et al. Elevated [18F]fluorodopamine turnover in brain of patients with schizophrenia: an [18F]fluorodopa/positron emission tomography study. The Journal of neuroscience: the official journal of the Society for Neuroscience. 2007;27(30):8080–7.

67. Howes OD, Bose SK, Turkheimer F, Valli I, Egerton A, Valmaggia LR, et al. Dopamine synthesis capacity before onset of psychosis: a prospective [18F]-DOPA PET imaging study. The American journal of psychiatry. 2011;168(12):1311–7.

68. Egerton A, Chaddock CA, Winton-Brown TT, Bloomfield MA, Bhattacharyya S, Allen P, et al. Presynaptic striatal dopamine dysfunction in people at ultra-high risk for psychosis: findings in a second cohort. Biological psychiatry. 2013;74(2):106–12.

69. Jauhar S, McCutcheon R, Borgan F, Veronese M, Nour M, Pepper F, et al. The relationship between cortical glutamate and striatal dopamine in first-episode psychosis: a cross-sectional multimodal PET and magnetic resonance spectroscopy imaging study. The lancet Psychiatry. 2018;5(10):816–23.

70. Martinot M, Bragulat V, Artiges E, Dolle F, Hinnen F, Jouvent R, et al. Decreased presynaptic dopamine function in the left caudate of depressed patients with affective flattening and psychomotor retardation. The American journal of psychiatry. 2001;158(2):314–6.

71. Bowden C, Cheetham SC, Lowther S, Katona CL, Crompton MR, Horton RW. Reduced dopamine turnover in the basal ganglia of depressed suicides. Brain research. 1997;769(1):135–40.

72. Dwivedi Y, Rizavi HS, Roberts RC, Conley RC, Tamminga CA, Pandey GN. Reduced activation and expression of ERK1/2 MAP kinase in the post-mortem brain of depressed suicide subjects. Journal of neurochemistry. 2001;77(3):916–28.

