## Supplementary Material for "Mesocorticolimbic circuit mechanisms underlying the effects of ketamine on dopamine: a translational imaging study"

**Supplementary information**

**Supplementary methods:**

All experiments were approved by the UK Home Office Animal (Scientific Procedures) Act (ASAP) 1986 and Regulation 7 of the Genetically Modified Organisms (Contained Use) Regulations 2000. All procedures were performed in accordance with the ASPA 1986 and EU directive 2010/63/EU as well as being approved by Imperial College Animal Welfare and Ethical Review Body. Mice were housed in individually ventilated cages and were maintained under a constant 12 h light-dark cycle (lights on at 7:00am) with access to food *ad libitum* (chow diet; RM3 SDS Essex UK Ltd) and water. Mice were group housed 2-5 mice per cage.

**Subjects**

The dopamine transporter *(DAT)::Cre* mice were maintained on C57BL/6 background. The mouse line expressing Cre recombinase under parvalbumin promoter/enhancer elements, referred to us parvalbumin (*PV)::Cre* was obtained from the Jackson Laboratory (008069) and maintained on a C57BL/6 background. C57BL/6 mice were obtained from Charles River Laboratories (Harlow, England).

**Genotyping**

The genotypes of both *DAT::Cre* and *PV::Cre* mice were confirmed by standard PCR amplification-based genotyping. 1µl of genomic DNA was added to 0.2 µM forward and reverse primers (DAT::Cre forward primer CCAGCTCAACATGCTGCACA, DAT::Cre reverse primer GCCACACCAGACACAGAGAT, PV::Cre wildtype forward primer CAGAGCAGGCATGGTGACTA, PV::Cre wildtype reverse primer AGTACCAAGCAGGCAGGAGA, PV::Cre mutant forward AAATGCTTCTGTCCGTTTGC, PV::Cre mutant reverse ATGTTTAGCTGGCCCAAATG) and 14 µl of PCR reagent mixture (Reddymix PCR Master Mix, ThermoScientific, USA). DAT::Cre genotyping reactions were carried out in a 15 µl final volume in a thermocycler (C100, BioRad) under the following PCR machine settings:95°C for 3 min, 5x(95°C for 30sec, 60°C for 30sec, 72°C for 1min), 25x(95 °C for 30 sec, 58°C for 30sec, 72°C for 1min), 72°C for 6 min, and then 12°C for 3min. PV::Cre PCR genotyping was carried out under the following conditions: 94°C for 3 min, 5x(94°C for 30sec, 64°C for 30sec, 72°C for 50sec), 5x(94 °C for 30 sec, 61°C for 30sec, 72°C for 50sec), 30x(94 °C for 30 sec, 56°C for 30sec, 72°C for 50sec), 72°C for 5 min, and then 12°C for 3min. All PCR products were separated by gel electrophoresis on a 3% agarose gel prepared with Tris acetate EDTA (ethylenediaminetetraacetic acid) (TAE) buffer (40mM Tris acetate pH 8.3, 1mM EDTA) and 0.02% ethidium bromide and were subsequently imaged by illumination with ultraviolet light.

**Experimental Design**

A *priori* power calculations were based on calculated or reported effect sizes in the literature and guided the choice of sample size for each experiment in order to maximise the likelihood of meaningful results and to simultaneously reduce the number of mice used in accordance with the refinement principle of the 3Rs (replacement, reduction, refinement) in pre-clinical research (https://www.nc3rs.org.uk/the-3rs). Mice were randomized to experimental treatment groups using the online random number generator (https://www.random.org/) and researchers performing the experiments were blinded to analysis and/or treatment unless otherwise stated.

**Drug injections**

Ketamine hydrochloride solid (Sigma Aldrich) was dissolved in 0.9% saline solution in 6mg/ml, injected at a volume of 5ml/kg body weight, thus administered at a dose of 30mg/kg (Figure 1). Entacapone solid (40mg/kg) (Sigma Aldrich) was dissolved in 10% dimethyl sulfoxide (DMSO) and 0.9% saline. Benserazide hydrochloride solid (10mg/kg) (Sigma Aldrich) was dissolved in de-ionised water. 25mg/ml Clozapine-N oxide stock solution was prepared by dissolving 1g of CNO (Key Organics) in 1ml of DMSO and 39ml of phosphate buffered saline (PBS’A’), at pH 7.2. CNO was further diluted in PBS ‘A’ to 0.02mg/ml. Clozapine-N-oxide was injected in *DAT::Cre* mice at a dose of 0.1mg/kg and in *PV::Cre* mice at a dose of 0.5mg/kg. For the stereotaxic surgery buprivacaine (2.5mg/ml) was applied subcutaneously at the sterilized incision site as an analgesic, while intraperitoneal carprofen injection (2.5mg/kg) was administered both during surgery and post-surgery in drinking water at a dose of 0.0272mg/ml from 50mg/ml stock for 2-4 days as an analgesic. SEP-0363856 (SEP856) was dissolved in 50mM Acetate buffer pH5.5 and administered at 3mg/kg and a volume of 2ml/kg in C57BL/6 mice. All drugs were i.p injected.

**PET analysis**

Emission data were histogrammed in 43 frames of increasing duration over the 120-min dynamic PET scan (frame intervals: 10 x 3s, 6 x 5s, 8 x 30s, 5 x 60s, 6 x 300s, 8 x 600s) and were then reconstructed into 3D images using a filtered back projection algorithm, correcting for CT attenuation, random, scatter and radiotracer decay.

**PET scanning region of interest definition**

PET images were analysed using the Inveon Research Workplace (IRW) software. For each scan, three regions of interest (ROIs) were drawn manually on summation radioactivity images at the level of the striatum (right and left) and cerebellum to extract time activity curves (TACs) (Supplementary figure 7) (Frame ranges that were summed= 2.5min-120min). Each ROI had ellipsoid shape (1, 2).

***K_i_ ^mod^* calculation**

Striatal dopamine synthesis capacity indexed by the rate of [^18^F]FDOPA uptake was calculated relative to the cerebellum [*K_i_ ^mod^* (min^-1^)]. To derive *K_i_ ^mod^* (min^-1^) extended Patlak graphical analysis was used. The Patlak method was adapted for reference tissue input function in place of the blood activity and accounting for loss of radiolabelled metabolites from trapped compartment by evaluating *k*_loss_ (min^-1^) according to Walker et al., 2013 (Equation 1), with the equilibrium time set to 20min (1-4).

Equation 1:

$$\frac{C_{Tissue}\left( t \right)}{C_{Reference} \left( t \right)}={K_{i}}^{mod}\theta\left( t \right)+V$$

With

$\theta\left( t \right)= \frac{\int_{0}^{t} C_{reference}\left( t^{'} \right)\exp\left( -k_{loss}\left( t-t^{'} \right) \right)dt'}{C_{reference}(t)}$

**Chemogenetics model**

Mice were anaesthetized with isoflurane and placed into a stereotaxic frame (David Kopf Instruments, Tujunga, CA, USA). Bilateral injections with 500 nL of Adeno-associated virus (AAV) construct carrying the hM4Di-mCherry or the hM3Dq-mCherry designer receptors exclusively activated by designer drugs (DREADD) receptors were administered at 150 nL/min using a Hamilton microinjection syringe.

**Virus construct and packaging**

Viruses used in this study were the AAV1-Ef1a-DIO-hM4Di-mCherry and the AAV1-Ef1a-DIO-hM3Dq and were generated as previously (5). The virus titres below 1.2x10^13^ genome copies (GC) ml-1 were used directly whilst the titres above this were diluted to 10^12^ genome copies (GC) ml-1 in sterile PBS A, with 5% glycerol (pH 7.2).

**Histology**

Mice were culled by cervical dislocation and brains were rapidly removed. Brains were post-fixed in 4% paraformaldehyde (PFA) overnight and then cryo-protected in 30% sucrose solution. Following sinking in sucrose, brains were rapidly frozen in isopentane (2-methylbutane) and stored in -80°C. Coronal sections of 70µm for VTA/SNc slices or 50µm for hippocampal/pre-limbic cortex slices were taken on a cryostat (Leica Biosystems).

**Immunohistochemistry (IHC)**

Standard IHC was used to label dopamine neurons or parvalbumin neurons and reporter expression was confirmed using confocal microscopy under the Leica SP confocal microscope (Leica Microsystems, UK). Day 1: Briefly coronal sections were washed four times (5min) in phosphate-buffered saline (PBS) on a shaker plate. Sections were subsequently blocked by incubating in PBS containing 0.2% Triton X-100 (0.2% PBS-TX) and 6% normal donkey serum (NDS) for 45 min at room temperature on a shaker plate. Sections were then incubated with primary antibody solution (2% NDS in 0.2% PBS-Triton X) (For tyrosine hydroxylase (TH): chicken anti-tyrosine hydroxylase antibody (1:1000, ABCAM ab76442, Cambridge, UK) and for parvalbumin (PV): rabbit anti-parvalbumin antibody (1:1000, Abcam ab11427) overnight at 4 °C on a shaker plate. Day 2: Following four (5min) washes in 0.2% PBS-Triton X, sections were incubated with secondary antibody solution (2% NDS in 0.2% PBS-Triton X) (for TH: Alexa Fluor 488-conjugated goat anti-chicken secondary antibody (1:1000, Life technologies; for PV: Alexa Fluor 488-conjugated goat anti-rabbit (1:1000), Invitrogen A11008) for 2 hours in the dark at room temperature on shaker plate. Subsequently sections were washed three times (5min) in 0.2% PBS-Triton X and twice (5min) in PBS. Sections were mounted in Vectorshield Mounting Medium (Vector Laboratories).

**Confocal microscopy and Image analysis**

Images were taken at a scan format of 512x512. Confocal settings including the number of z-stacks and the z-step size were kept constant. Slices were evaluated for fluorescence under settings for 568, 488 and 400nm emissions. Each cortical slice was imaged across the PLc region between bregma +1.3 and +2.0 (three images per slice). Each hippocampal slice was imaged across the vSub between bregmas -3.40 and -3.64. Six slices per each of the two sub-regions of interest were analysed per subject. For each slice a z-stack of 8 images was obtained (corresponding to 1.4 µm on the z-axis). All PV neurons in the images were analysed for PV and DAPI content. The median fluorescence/cell was averaged across all imaged slices of the subject and the mean fluorescence intensity/cell/subject was then expressed as percent of control conditions as previously reported (6).

**In vitro electrophysiology**

Patch-clamp recordings in brain slices were carried out as described previously (5). Briefly, *PV::Cre* mice were stereotaxically injected with hM3Dq and two weeks later brains were rapidly transferred to an ice-cold artificial solution including 2.5 mM KCl, 1.25 mM NaH_2_PO_4_,28mM NaHCO_3_, 0.5 mM CaCl_2_, 7 mM MgCl_2_, 7 mM D-glucose, and 235 mM sucrose and they were equilibrated with 95% O_2_, 5% CO_2_ (pH 7.4). 350µm coronal brain slices were prepared using a Vibratome Series 1000. Ventral hippocampal slices were maintained at room temperature in an external solution containing 125 mM NaCl, 2.5 mM KCl, 1.25 mM NaH_2_P_4_, 25 mM NaHCO_3_,2 mM CaCl_2_, 1 mM MgCl_2_, 10 mM D-glucose, 15 mM D-mannitol, equilibrated with 95% O_2_, 5% CO_2_ (pH 7.4). PV neurons were visualized using epifluorescence using an upright Slicescope (Scientifica) microscope equipped with X40 objective. Changes in membrane potential were recorded. Application of CNO was performed through the bath perfusion system at the concentration indicated.

**Supplementary Figures:**

**Supplementary Figure 1: Sub-chronic ketamine administration induces locomotor hyperactivity and sensitization as measured using the open field test.** (a) Timeline and experimental design. Mice received either saline (*n*=14) or ketamine (*n*=14) once daily for five consecutive days and all mice received ketamine on day 7. (b-d) Distance travelled (m) in 5 min bins on days 1, 5 and 7. (e) Total distance travelled (m) in 30min post drug administration expressed as a percentage of saline. Saline (*n*=14) and Ketamine (*n=14*). Sub-chronic ketamine induced locomotor sensitization. (f) Representative trajectories from saline- and ketamine-treated mice on days 1, 5 and 7. Data represent mean ± S.E.M. ****P<0.001, *P<0.05*

**Supplementary Figure 2: Inhibition of midbrain dopamine neuron activity prevents the sub-chronic ketamine-induced effects on locomotor sensitization.** (a) Timeline and experimental design. Mice received CNO or saline 30min prior to ketamine or saline administration for 5 days and locomotor activity was measured in the open field test. (b+c) Distance travelled (m) in 5 min bins on days 1 and 5. (d) Representative trajectories of mice from each of the four experimental groups (Sal/Sal, CNO/Sal, Sal/Ket, CNO/Ket) on day 5. Data represent mean ± SEM. Sal/Sal *n*= 10, CNO/Sal *n*= 9, Sal/Ket *n* = 18, CNO/Ket *n*= 19

**Supplementary Figure 3: Parvalbumin (PV) interneuron activation reduces the effects of sub-chronic ketamine on locomotor activity.** (a) Timeline and experimental design. Mice received CNO or saline 30min prior to ketamine or saline administration for 5 days and locomotor activity was measured. (b+c) Distance travelled (m) in 5 min bins on days 1 and 5. (d) Representative trajectories of mice from each of the four experimental groups on day 5. Data represent mean ± SEM. Sal/Sal *n= 7*, CNO/Sal *n= 9*, Sal/Ket *n= 16*, CNO/Ket *n= 16*

**Supplementary Figure 4: *In vivo* CNO administration does not prevent the effects of ketamine on striatal dopamine synthesis capacity and locomotor sensitization**. (a+b) Distance travelled (m) in 5 min bins on days 1 and 5. (c) Total distance travelled (m) in 30min post drug administration. Sub-chronic ketamine treatment induced locomotor sensitization in both groups (*n=6* per group). (d) Striatal dopamine synthesis capacity (*K*_i_ *^mod^* /minute), in Sal/Ket and CNO/Ket groups in *DAT::Cre* mice that had received control virus injections. ns- not significant. Data represent mean ± SEM.

**Supplementary Figure 5:** No expression of mCherry in PV or TH immunolabelled neurons in wild type mice two weeks following stereotaxic surgery of (a) hM3Dq, scale bar=50µm or (b) hM4Di, scale bar=50µm constructs.

**Supplementary Figure 6:** Time activity curves show mean striatal and cerebellar [18F]-FDOPA radioactivity signal during the 2hour PET scan duration. The radioactivity is presented as standardized uptake values (SUVs), corrected for mouse body weight, injected dose and time of injection of the radiotracer. (a) C57BL/6 wild type PET experiment, (b) *DAT:Cre* PET experiment, (c) *PV:Cre* PET experiment.

**Supplementary Figure 7:** Representative images showing the delineation of the striatal and cerebellar regions of interest. The image depicts higher striatal [^18^F]-DOPA uptake.

**Supplementary Table 1:** Summary of scan characteristics and scan parameters for the C57BL/6 PET experiment. Abbreviations: ^a^ Independent-samples t-tests, ns - non significant, ****P<0.001*

| Sample characteristic and scan parameters | Ketamine treated [mean(SEM)] | Controls  [mean(SEM)] | Group comparisons  *t_df_* p^a^ |
| --- | --- | --- | --- |
| Weight (g) | 24.2 ± 0.93 | 24.8 ± 0.37 | t_14_=.68, *P=0.51*, ns |
| Injected dose (MBq) | 3.49 ± 0.72 | 4.45 ± 1.05 | t_14_=.76, *P=0.46*, ns |
| Specific activity (MBq/µmol) | 0.06 ± 0.02 | 0.06 ± 0.01 | t_10_=-0.06, *P=0.95*, ns |
| *K_i_*^mod^ (min^-1^) | 0.044 ± 0.005 | 0.015 ± 0.002 | t_13_= 4.74, ****P<0.001* |
| *K_i_*^Cer^ (min^-1^) | 0.011 ± 0.001 | 0.010 ± 0.001 | t_14_= 0.66, *P=0.52* |
| *k* _loss_ (min^-1^) | 0.043 ± 0.009 | 0.020 ± 0.007 | t_14_= 0.33, *P=0.75* |

**Supplementary Table 2:** Summary of scan characteristics and scan parameters for the *DAT::Cre* PET experiment. ***P<0.01, *P<0.05*

| **Parameter** | **Dopamine neuron firing manipulation**  **CNO vs Saline** | | **Ketamine vs Saline treatment** | | **Interaction between dopamine neuron firing manipulation and ketamine vs saline treatment** | |
| --- | --- | --- | --- | --- | --- | --- |
|  | **F** | **p** | **F** | **p** | **F** | **p** |
| *K*_i_^mod^ (min^-1^) | F_1,44_=10.82 | 0.002** | F_1,44_=3.012 | 0.08 | F_1,44_= 12.51 | 0.001** |
| *K*_i_^cer^ (min^-1^) | F_1,46_=0.26 | 0.61 | F_1,46_=1.84 | 0.18 | F_1,46_=10.84 | 0.002** |
| k_loss_ (min^-1^) | F_1,46_=5.27 | 0.03* | F_1,46_=4.29 | 0.04* | F_1,46_=3.45 | 0.07 |
| Weight (g) | F_1,46_=0.40 | 0.53 | F_1,46_=1.10 | 0.30 | F_1,46_=0.21 | 0.65 |
| Injected dose (MBq) | F_1,46_=0.32 | 0.57 | F_1,46_=4.93 | 0.03* | F_1,46_=0.11 | 0.74 |
| Specific Activity (MBq/µmol) | F_1,46_=0.35 | 0.56 | F_1,46_=11.9 | 0.001** | F_1,46_=1.81 | 0.19 |

**Supplementary Table 3:** Summary of scan characteristics and scan parameters for the *PV::Cre* PET experiment. Abbreviations: ^a^ Independent-samples t-tests, ns - non significant, ***P<0.01*

| Sample characteristic and scan parameters | Sal/Ket (n=11)  [mean(SEM)] | CNO/Ket (n=10)  [mean(SEM)] | Group comparisons  *t_df_* p^a^ |
| --- | --- | --- | --- |
| Weight (g) | 27.74 ±0.76 | 28.44±0.48 | t_19_=0.77, *P=0.45*, ns |
| Injected dose (MBq) | 4.62 ±0.68 | 4.18±0.38 | t_19_=0.54*, P=0.59*, ns |
| Specific activity (MBq/µmol) | 0.02 ± 0.002 | 0.02 ± 0.002 | t_14_=0.5 *, P=0.51*, ns |
| *K_i_*^mod^ (min^-1^) | 0.033 ± 0.003 | 0.023 ± 0.002 | t_19_=3.51, ***P<0.01* |
| *K_i_*^Cer^ (min^-1^) | 0.011 ± 0.0005 | 0.011 ± 0.0003 | t_19_=0.90, *P=0.38*, ns |
| *k* _loss_ (min^-1^) | 0.027 ± 0.002 | 0.017 ± 0.002 | t_19_=3.65, ***P<0.01* |

**References**

1.	Walker MD, Dinelle K, Kornelsen R, Lee A, Farrer MJ, Stoessl AJ, et al. Measuring dopaminergic function in the 6-OHDA-lesioned rat: a comparison of PET and microdialysis. EJNMMI research. 2013;3(1):69.

2.	Bonsall DR, Kokkinou M, Veronese M, Coello C, Wells LA, Howes OD. Single cocaine exposure does not alter striatal presynaptic dopamine function in mice: an [18 F]-FDOPA PET study. Journal of neurochemistry. 2017.

3.	Holden JE, Doudet D, Endres CJ, Chan GL, Morrison KS, Vingerhoets FJ, et al. Graphical analysis of 6-fluoro-L-dopa trapping: effect of inhibition of catechol-O-methyltransferase. Journal of nuclear medicine : official publication, Society of Nuclear Medicine. 1997;38(10):1568-74.

4.	Patlak CS, Blasberg RG. Graphical evaluation of blood-to-brain transfer constants from multiple-time uptake data. Generalizations. Journal of cerebral blood flow and metabolism : official journal of the International Society of Cerebral Blood Flow and Metabolism. 1985;5(4):584-90.

5.	Viskaitis P, Irvine EE, Smith MA, Choudhury AI, Alvarez-Curto E, Glegola JA, et al. Modulation of SF1 Neuron Activity Coordinately Regulates Both Feeding Behavior and Associated Emotional States. Cell reports. 2017;21(12):3559-72.

6.	Behrens MM, Ali SS, Dao DN, Lucero J, Shekhtman G, Quick KL, et al. Ketamine-induced loss of phenotype of fast-spiking interneurons is mediated by NADPH-oxidase. Science. 2007;318(5856):1645-7.
