## Supplementary figures and images for "Mesocorticolimbic circuit mechanisms underlying the effects of ketamine on dopamine: a translational imaging study"

### Supplementary Figure 1

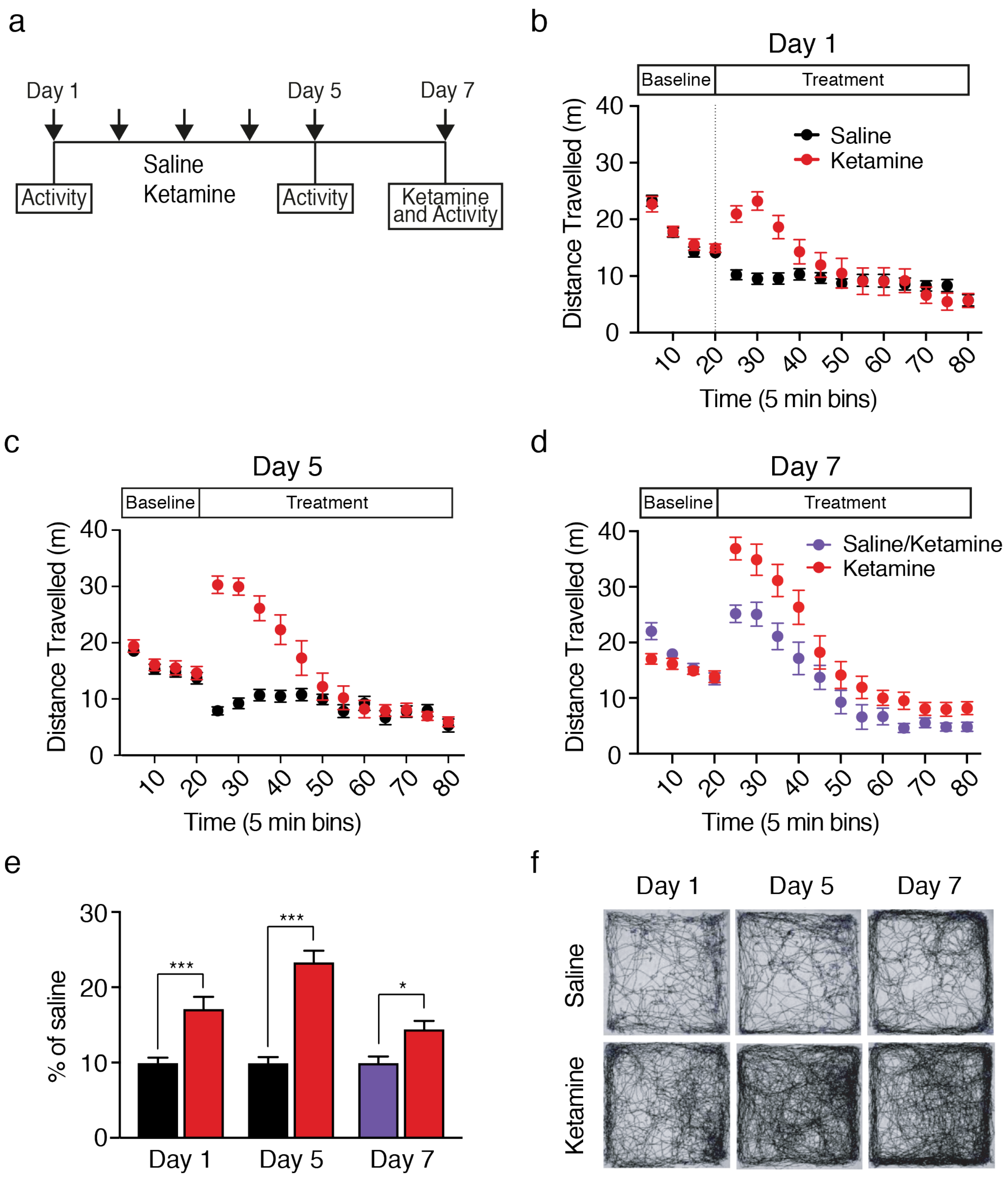

### Supplementary Figure 2

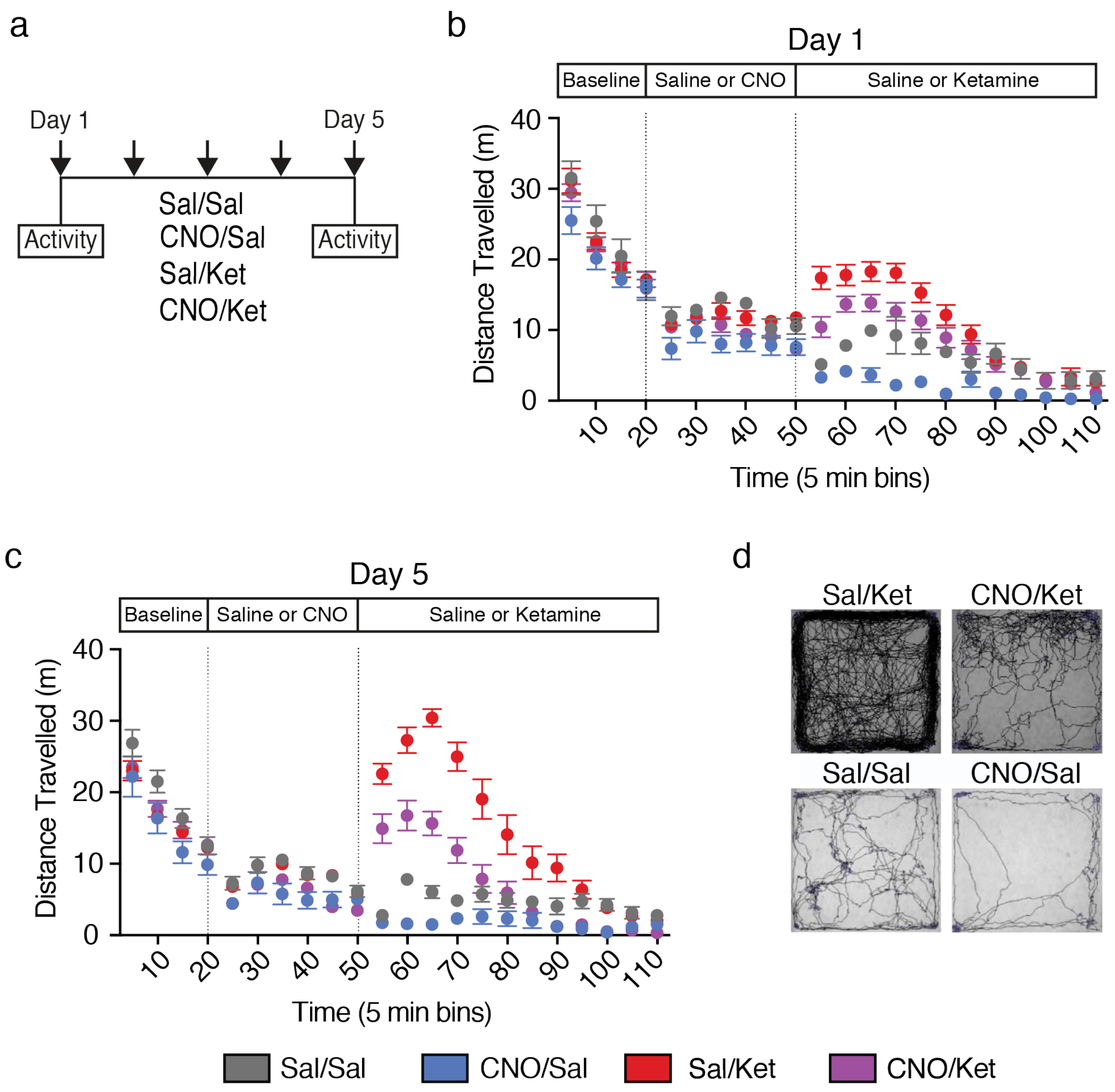

### Supplementary Figure 3

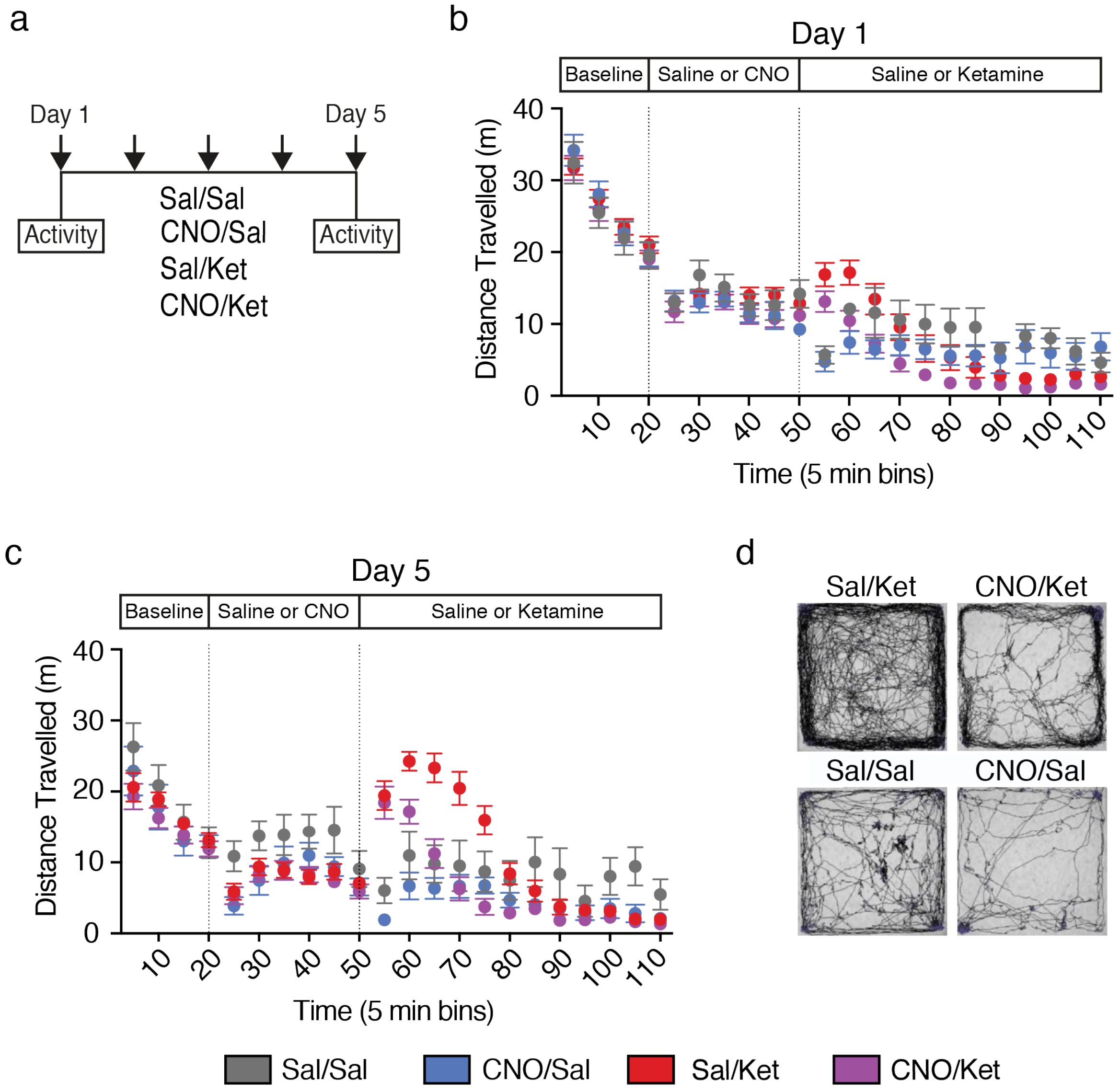

### Supplementary Figure 4

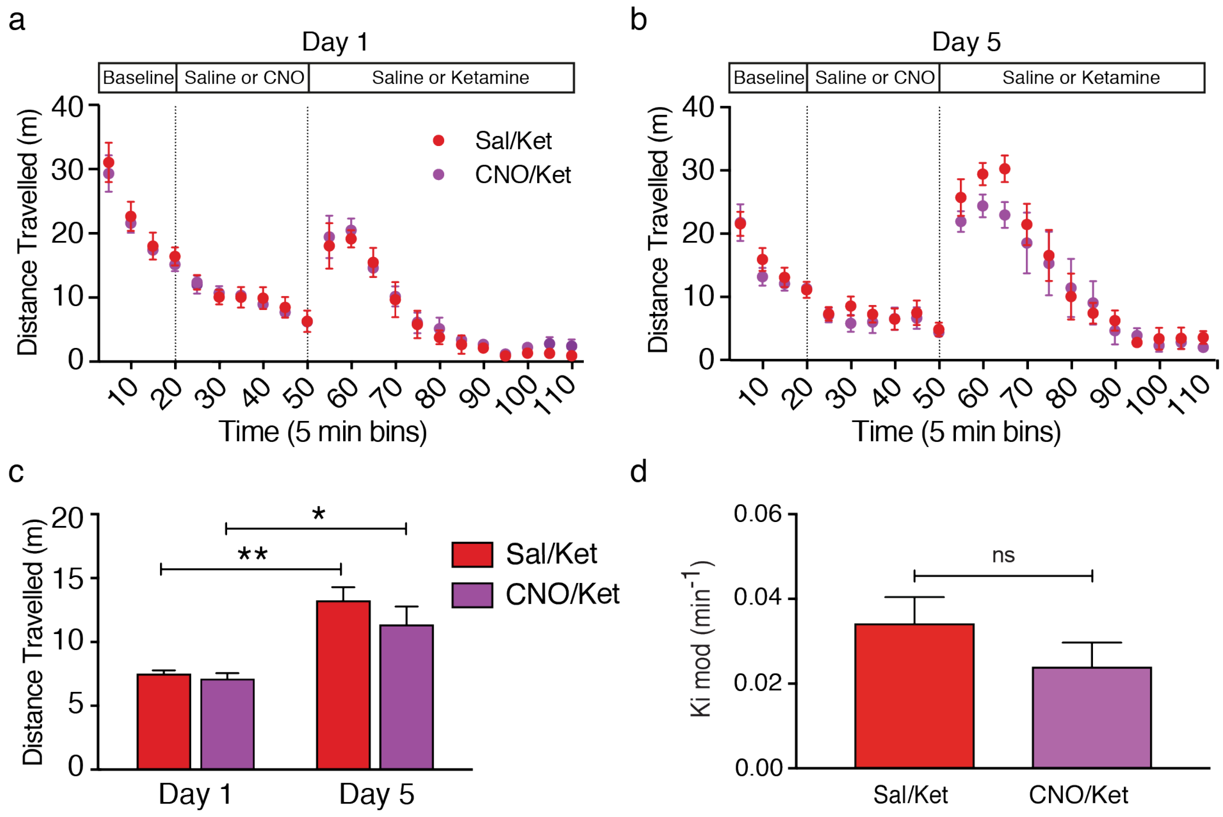

### Supplementary Figure 5

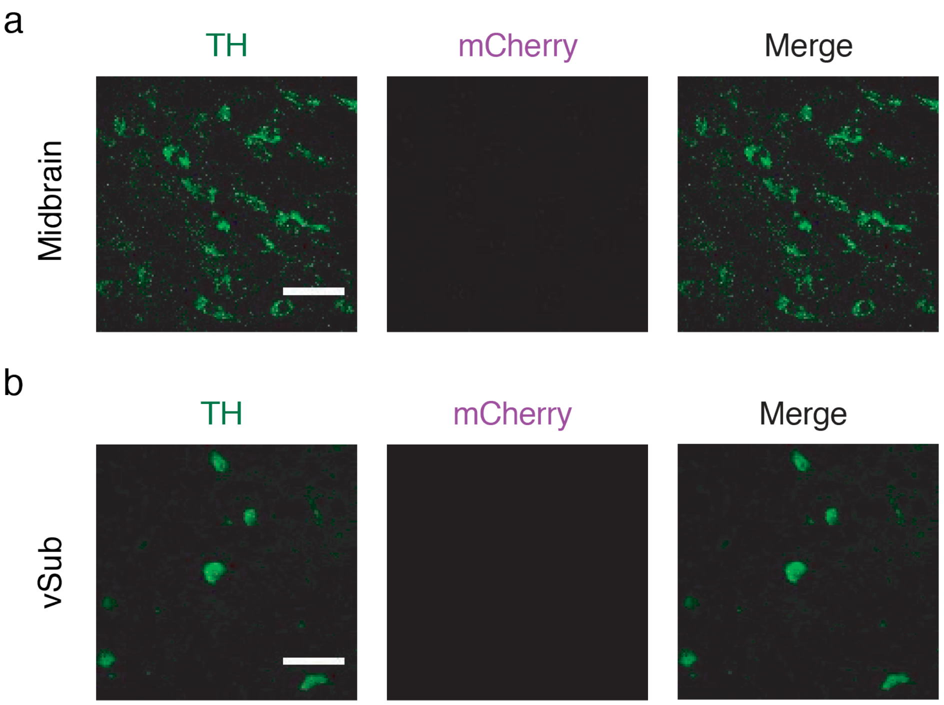

### Supplementary Figure 6

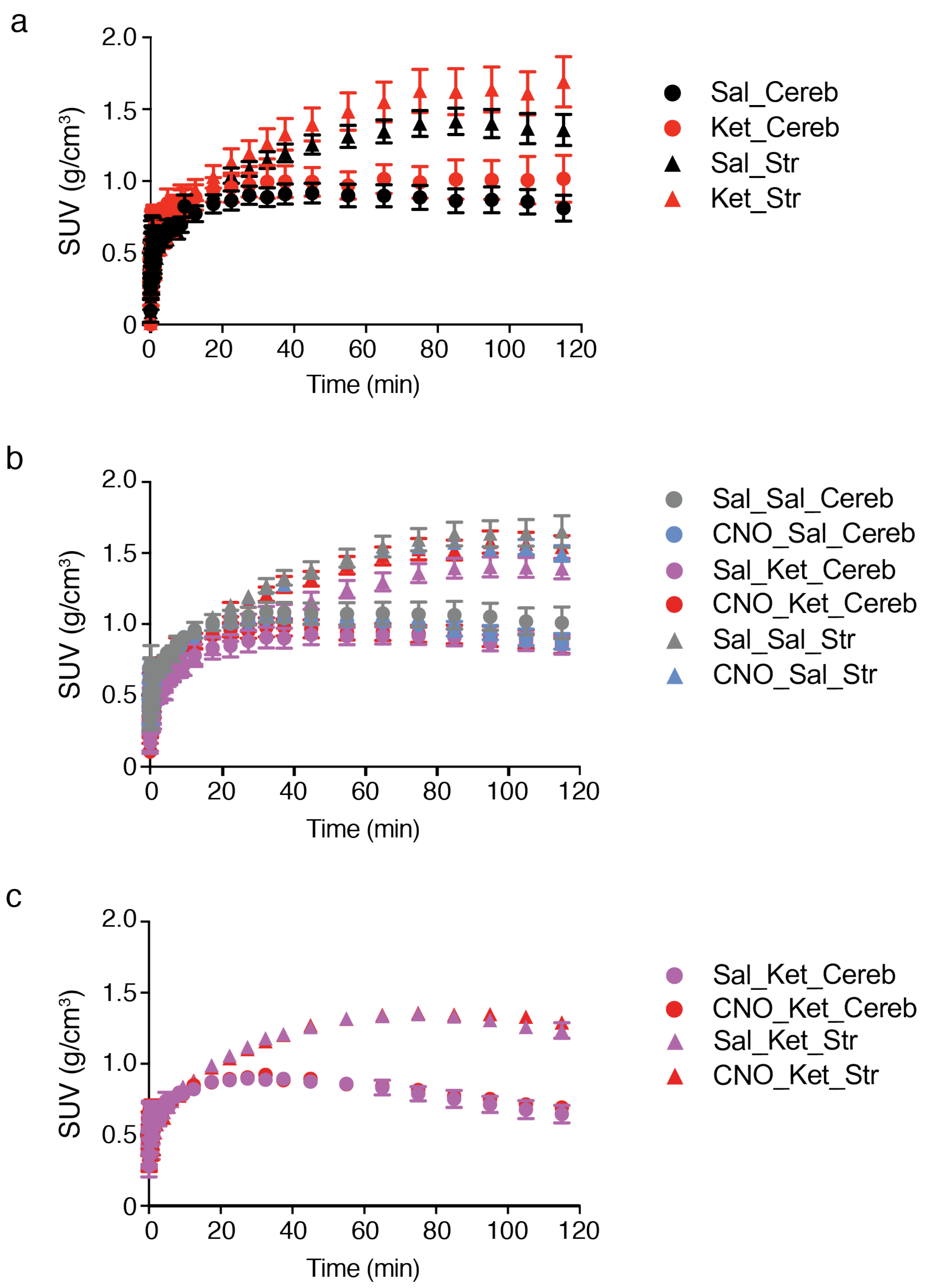

### Supplementary Figure 7

a

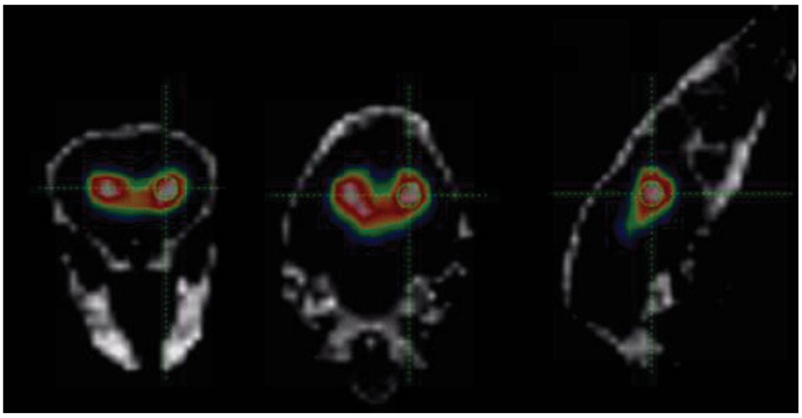

b

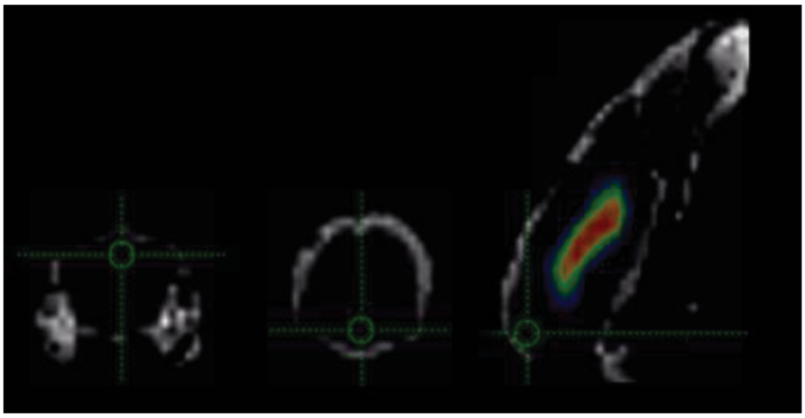
